# Targeting aggressive B-cell lymphomas through pharmacological activation of the mitochondrial protease OMA1

**DOI:** 10.1101/2022.06.12.495213

**Authors:** Adrian Schwarzer, Matheus Oliveira, Marc-Jens Kleppa, Scott D. Slattery, Andy Anantha, Alan Cooper, Mark Hannink, Axel Schambach, Anneke Dörrie, Alexey Kotlyarov, Matthias Gaestel, Todd Hembrough, Jedd Levine, Michael Luther, Michael Stocum, Linsey Stiles, David Weinstock, Marc Liesa, Matthew J. Kostura

## Abstract

Constitutive activation of the ATF4-mediated integrated stress response (ATF4-ISR) is common in cancer and buffers the metabolic challenges imposed by rapid proliferation. However, hyperactivation of the ISR can induce apoptosis. Here we demonstrate that novel pyrazolo-thiazole derivates activate the mitochondrial protease OMA1 which subsequently induces apoptosis in diffuse large B-cell lymphoma (DLBCL) cells. Apoptosis is dependent on the OMA1 mediated cleavage of DELE1 which leads to activation of HRI and induction of the ATF4 ISR. Screening in 406 cancer cell lines identified an inverse correlation between sensitivity to OMA1 activators and expression of the mitochondrial protein FAM210B. Ectopic overexpression of FAM210B specifically blocks OMA1 activation and apoptosis induction by pyrazolo-thiazole activators in DLBCL. OMA1 activators, including the preclinical candidate BTM-3566, selectively killed ABC, GCB, and double-hit DLBCL lines and induced complete tumor regression across a panel of DLBCL patient-derived xenografts.

**Significance:** Here we describe a novel class of small molecules that activate the mitochondrial protease OMA1 and induce therapeutic responses in DLBCL preclinical models in vitro and in vivo. OMA1 activation drives apoptosis through ATF4-ISR, an orthogonal mechanism to current therapies.

## INTRODUCTION

Malignant cells and tumors require robust homeostatic regulation to adjust the metabolic demands of rapid proliferation with nutrient availability and accumulation of metabolic end products to survival (1). To meet these demands, tumors cells rely on endogenous stress survival pathways(2). On the other hand, the utilization of stress signaling pathways also creates dependencies that present therapeutic opportunities.

Several Mitochondria Quality Control (MQC) pathways serve to maintain the integrity of mitochondria. MQC pathways sense changes in mitochondrial oxidative phosphorylation, membrane potential, proteostasis and translation of mitochondrial DNA encoded proteins (3–5). MQC pathways signal to cytosolic pathways that control the ATF4-Integrated Stress Response (ISR), an adaptive gene expression program controlling amino acid biosynthesis and transport, redox homeostasis and enhanced protein folding (3,6,7). The ATF4-ISR facilitates adaptation to the tumor microenvironment and can be pharmacologically targeted (8,9). At the same time, persistent ISR activation can sensitize cells to apoptotic stimuli by altering the levels of pro- and anti-apoptotic proteins (10–15).

Herein we describe the preclinical pharmacology of a novel class of pyrazolo-thiazole derivatives (hereafter BTM-compounds), first described for their ability to induce cell growth arrest and cell death in a wide variety of solid and hematopoietic tumor cell lines (16). We demonstrate that select BTM-compounds specifically hyperactivate the ISR through activation of the MQC protease OMA1, thereby shifting the equilibrium from ATF4-ISR controlled pro-survival effects towards tumor cell death. Sensitivity to BTM-compounds is suppressed by the mitochondrial protein FAM210B, which is minimally expressed across multiple subtypes of diffuse large B-cell lymphoma, providing a therapeutic opportunity for OMA1 activators.

## RESULTS

### BTM-compounds induce DLBCL cell death *in vitro*

We previously described a series of pyrazolo-thiazole compounds that displayed anti-proliferative activity across a range of hematopoietic and solid tumor cell lines (16). We sought to further improve this class of compounds by optimizing the substituents around the pyrazolo-thiazole core. The medicinal chemistry approach was guided by the potency of derivatives to induce cell death in human lymphoma cell lines and spare non-transformed B-cells. This effort yielded two lead compounds, BTM-3528 and BTM-3566, with good solubility and potent *in vitro* activity, together with BTM-3532, a structurally related but biologically inactive compound (**Fig. 1A**). We assessed the anti-proliferative activity of BTM-3528 and BTM-3566 in a diverse panel of 99 tumor cell lines, including 19 hematopoietic lines and 80 solid tumor lines. BTM-3528 and BTM-3566 demonstrated equivalent activity across solid tumor cell lines, with the greatest activity observed in lung, colorectal and pancreatic cancer and lack of activity in skin and breast cancer cell lines **(Fig. 1B and C; Supplementary Table S1)**. In contrast, hematopoietic tumor lines were broadly responsive to both BTM compounds (**Fig. 1B and C; Supplementary Table S1**), with the most sensitive being Burkitt lymphoma and DLBCL lines **(Fig. 1C, Supplementary Table S1)**. A comparison of BTM-3528 and BTM-3566 revealed potent activity against DLBCL lines of diverse genotypes, including *MYC-*rearranged (“double” and “triple” hit) lymphomas, with >90% growth inhibition and IC_50_ values of 0.16-0.57 μM (**Supplementary Table S2**). Cell viability experiments showed that BTM-3528 and BTM-3566 induced rapid apoptosis in lymphoma lines, with nearly complete killing at 24 hrs (**Fig. 1D, Supplementary Fig. S1A**), whereas the inactive compound BTM-3532 had no effect on survival or proliferation (**Supplementary Fig. S1B, D-E)**. Apoptosis induced by BTM-3528 and BTM-3566 was accompanied by caspase 3/7 activation in B-cell lymphoma lines. This result contrasted with solid tumor lines, which exhibited cell growth inhibition due to G1 arrest, but no apoptosis or caspase activation (**Fig. 1E, Supplementary Fig. S1C-E**).

**Fig 1.**
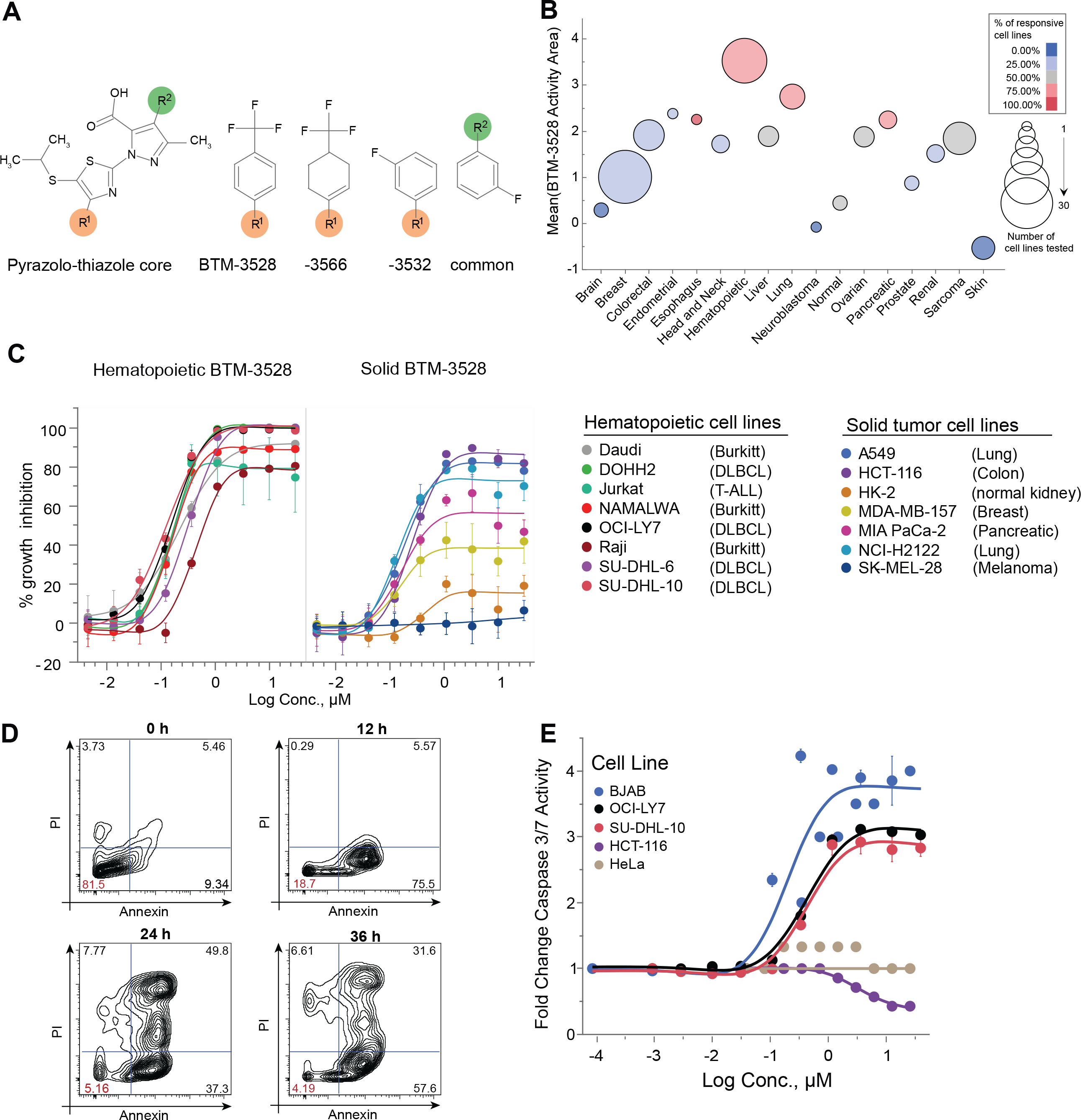
Pyrazolo-thiazole derivates are novel anti-cancer agents. **A)** Chemical structures of the BTM pyrazolo-thiazole series. BTM-3528 and BTM-3566 are active versions of the series. BTM-3532 is a closely related inactive member of the series. **B)** Summary of activity of BTM-3528 in 99 tumor cell lines from different tumor entities plotted as Mean Activity Area (MAA, integrated potency and magnitude of cell growth inhibition, in the CellTiterGlo assay, see materials and methods) over tumor tissue of origin. **C)** Dose response curves of select hematopoietic tumor cell lines (left panel) and solid tumor lines (right panel) treated with BTM-3528**. D)** Annexin-PI staining performed in BJAB cells incubated with 2μM BTM3528 for 12, 24, and 36 hrs. The percentage of surviving cells is indicated in red font in the lower left quadrant. **E)** Induction of Caspase 3/7 activity in selected DLBCL and solid tumor cell. All data are plotted as the mean (+/− SD, n=3) as compared to vehicle.

### BTM-3566 has favorable pharmacokinetic properties and potent *in vivo* activity in human cell line and patient-derived xenograft models

To investigate the pharmacokinetic properties of BTM-3566, we performed i.v./p.o. crossover studies in mice (**Fig 2A-C**). Bioavailability was > 90% and the terminal half-life of 4.4 to 6.6 hours was acceptable for once-daily oral dosing.

**Fig 2.**
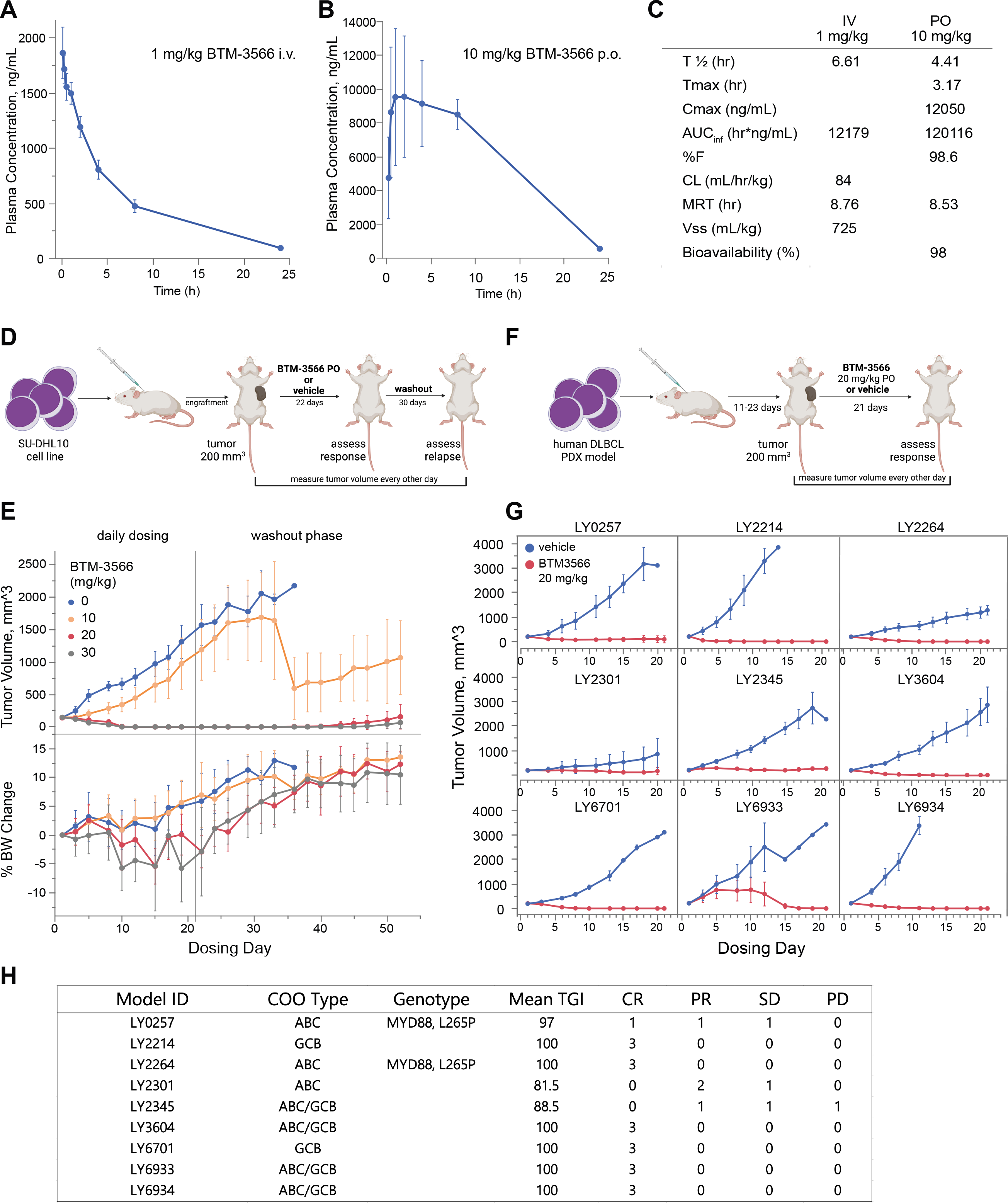
BTM-3566 has favorable pharmacokinetics and induces durable remissions in DLBCL-xenograft models. **A)** Plasma concentration of BTM-3566 after bolus iv administration of 1 mg/kg BTM-3566. N= 3 animals per condition. **B)** Plasma concentration of BTM-3566 after oral gavage dosing of 10 mg/kg BTM-3566 **C)** PK parameters for dosing of BTM-3566 in the mouse. **D)** Graphical representation of the Su-DHL-10 xenograft model: Cells were implanted subcutaneously in the flank of SCID beige (C.B-17/IcrHsd-Prkdc ^scid^Lyst^bg-J^) mice and grown until tumor volume reached 200 mm^3^ prior to randomization into four groups. Mice were then dosed po, qd with vehicle or a solution of 10, 20 or 30 mg/kg BTM-3566 dissolved in 5%NMP/15% Solutol/10% PEG400/70% D5W. **E)** SU-DHL-10 xenograft model: Upper panel: tumor volume over time, Bottom Panel: body weight over time. The vertical line at day 21 represents the end of drug dosing. All data are the mean +/− SD (n=10 animals). **F)** Graphical representation of the DLBCL PDX-models: nine human DLBCL PDX tumors were established in SCID mice and grown for 11-23 days to reach 200 mm^3^ before randomization. All groups received drug or vehicle for 21 days. Treatment arms received BTM-3566, 20 mg/kg, PO, qd. Tumor volume was measured and recorded every other day. **G)** Tumor growth curves over time in nine PDX models. All data represented as mean ± SD (n=3 animals). •=Vehicle; •= BTM-3566. **H)** Tabulated results of PDX model testing. CR, Complete Response, no palpable tumor; PR, Partial Response, palpable tumor with volume >50% less than baseline; SD, Stable Disease, tumor volume is not increased above baseline; PD, Progressing Disease where tumor is larger than starting tumor volume.

Next, we assessed the therapeutic activity and potency of BTM-3566 in a human xenograft model using the double-hit lymphoma DLBCL tumor line SU-DHL-10 (**Fig 2D**). At the 10 mg/kg dose, we observed delayed tumor growth that was lost with further dosing (**Fig. 2E upper panel**). At doses at or above 20 mg/kg daily, BTM-3566 treatment resulted in complete responses (CR, defined as no palpable tumor) in all animals by ten days of dosing and maintained for 21 days of dosing. To assess whether BTM treatment would induce durable responses, animals were followed for thirty additional days after cessation of dosing. Thirty-day tumor-free survival was maintained in 40% of animals dosed with 20 mg/kg and 60% of animals dosed with 30 mg/kg BTM-3566 (**Fig. 2E upper panel**). Body weight loss was dose-dependent but <10% at the 20 mpk dose level (**Fig. 2E lower panel**). In the 30 mg/kg dose group, two of 10 mice exceeded 20% body weight loss, necessitating an unscheduled dose holiday. Weight loss in both groups was reversible with cessation of dosing. (**Fig. 2E lower panel**).

Having established the 20 mg/kg dose as effective and well tolerated in the SU-DHL10 model, we next tested this dose of BTM-3566 in 9 human DLBCL patient-derived xenograft (PDX) models representing ABC- and GCB DLBCL-subtypes and including high-risk genotypes (**Fig. 2F-H**). Treatment with 20 mg/kg of BTM-3566 resulted in CR in all 3 mice from 6 of 9 PDX models. When pooling the 27 mice in the treatment arms of the nine models, CR was observed in 66% (19 of 27), partial response (PR) occurred in another four mice, 2 mice had stable tumors, and two had progressive disease. In summary, the single-agent overall response rate (CR+PR) was 85.2 % with all models having at least one animal exhibiting full or partial regression. (**Fig. 2H**).

### BTM-3566 and BMT-3528 induce activation of the ATF4-linked ISR

To elucidate the mechanism of action of the active BTM compounds, we next examined the transcriptional changes induced by compound treatment. We chose the colon cancer cell line HCT-116 for this experiment because all DLBCL cell lines rapidly underwent apoptosis upon exposure to BTM compounds, preventing time-course experiments. We performed RNA sequencing (RNA-seq) in HCT-116 cells treated with BTM-3528 at 10, 1, or 0.1 μM for 1, 2, 4, 6, or 8 hours, and vehicle-treated cells. Treatment with BTM-3528 induced rapid changes in gene expression, with the first differentially expressed genes appearing at the 4-hour time point (**Fig 3A**; **Supplemental Figure S2 A, B**). Gene set enrichment analysis revealed upregulation of genes associated with the ATF4-linked integrated stress response (**Supplemental Figure S2C**). Expression of *ATF4* itself was upregulated by BTM-3528 treatment and many of the most upregulated genes at each time point were direct ATF-4 target genes (**Fig. 3A,** in red font).

**Fig. 3.**
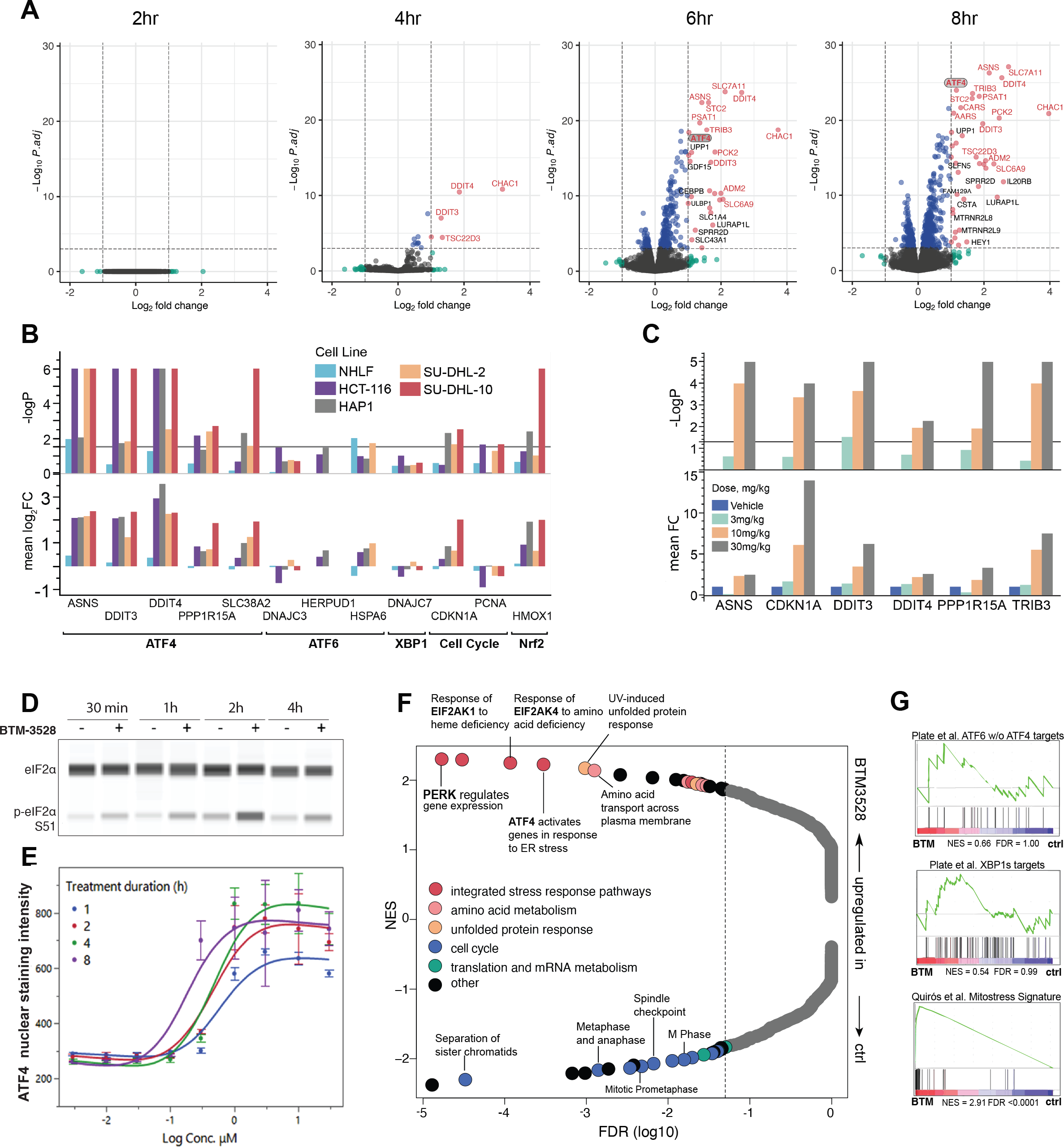
BTM-3528 activates the ATF4-linked mitochondrial stress response. **A)** Volcano plots showing the adjusted P-value (−log_10_ P) versus the fold change (log2) for 14,287 genes after 2,4,6, and 8 hrs of treatment with BTM-3528 versus vehicle. The dashed lines indicate the log2-FC cutoff =1.0 and the adj.p-value cutoff = 10^−3^. ATF4-target genes (71) are indicated by red font. **B)** qPCR analysis of a panel of cell lines treated with BTM-3528 for 8 hours. Lower panel shows gene changes expressed as mean log_2_-FC compared to vehicle (n=3). Upper panel shows the corresponding -LogP value for each triplicate sample as compared to vehicle. **C)** BTM-3566 induces tumor ATF4 gene expression in a dose dependent manner *in vivo*. On day 5 of dosing tumor tissue was harvested for qPCR analysis of mRNA expression. All data are expressed as the mean fold-change (lower panel) and corresponding -LogP (upper panel) for the contrast BTM vs vehicle for each gene. **D)** Immunoblot of phosphorylated eIF2α in HCT-116 following treatment with BTM-3528 for the indicated time. **E)** Dose response of nuclear ATF4 protein abundance in HCT-116 cells treated with BTM3528 for the indicated time and dose. **F)** Gene Set Enrichment Analysis (GSEA) of 1773 canonical pathway gene sets from MSigDB C2 of BTM-3528 treated samples versus controls at 8 hrs. Plotted are normalized enrichment scores (NES) against the false discovery rate (FDR). The enrichment cutoff (FDR < 0.05) is indicated by the dashed line. **G)** Upper Panel: Gene set enrichment analysis for ATF6 target genes; Middle Panel XBP1 target genes (26); lower panel, mitostress signature (8) in HCT116 cells treated with BTM3528 for 8 hrs.

Using qPCR we confirmed that BTM-3528 increased the expression of ATF4 target genes in responsive solid and hematopoietic tumor lines including HCT116, the chronic myelogenous leukemia cell line HAP1, and two DLBCL cell lines SU-DHL-2 and SU-DHL-10 **(Fig 3B)**. In contrast, ATF4-regulated genes were not induced in normal human lung fibroblasts. Consistent with the RNASeq-data, transcriptional responses to BTM-3528 were selective for ATF4-but not ATF6- or ERN1/XBP1-regulated transcripts associated with ER stress **(Fig. 3B**). Finally, we tested the ability of BTM-3566 to induce the eIF2α-ATF4-ISR *in vivo* in the SU-DHL-10 xenograft model of DLBCL. Mice were dosed with 3, 10, and 30 mg/kg per day of BTM-3566 for 5 days. Six hours following drug administration, the expression of ATF4 target genes (*ASNS*, *DDIT3*, *TRIB3*, and *CDKN1A*) in tumors increased in a dose-dependent manner **(Fig. 3C)**. Both the reduction in tumor volume and the increase in ATF4 target gene expression for *ASNS*, *CDKN1A*, *TRIB3*, and *DDIT3* correlated with overall BTM-3566 exposure in the mouse (**Supplementary Fig. S4)**.

To confirm activation of the ATF4 regulated ISR, we determined whether phosphorylation of eIF2α and nuclear localization of ATF4 protein were increased in HCT-116 and BJAB DLBCL cells treated with BTM-3528. Western blotting demonstrated a time-dependent increase in eIF2α phosphorylation (**Fig. 3D**), while immunofluorescent staining of nuclear ATF4 revealed a dose- and time-dependent accumulation of ATF4 protein in the nuclei of HCT-116 cells treated with BTM-3528 (**Fig. 3E and Supplementary Fig. S3**). Induction of ATF4 by BTM-3528 occurred within 1 hour following treatment with an EC_50_ of 0.19 μM, consistent with the early upregulation of ATF4 target genes observed in our RNASeq data.

ATF4 regulated gene expression is controlled by phosphorylation of the translation initiation factor eIF2α (17–19). Gene set enrichment implicated three kinases, HRI, GCN2, and PERK as potential drivers of BTM-3528 mediated ATF4 upregulation (**Fig. 3F**): HRI (EIF2AK1) is activated by heme-deprivation, protein misfolding and mitochondrial stress (20). PERK (EIF2AK3) is activated by the unfolded protein response and ER stress (21,22), and GCN2 (EIF2AK4) is activated by nutritional stress, specifically amino acid deprivation leading to accumulation of deacylated tRNA (9,23). Since ER-stress also activates the transcription factors ATF6 and XBP1S independently of the activation of PERK, we determined whether ATF6 and XBP1 target genes were affected by BTM-3528 treatment. We found that genes regulated by ER-stress and the transcription factors ATF6 or XBP1 (24) were not upregulated by treatment with BTM-3528 (**Fig. 3G, upper and middle panel**) suggesting that PERK was not activated by BTM-3528. Using a similar methodology, we found that the transcriptional changes induced by BTM-3528 treatment were highly similar to those observed in the context of mitochondrial stress (**Fig. 3G, lower panel**) (6). Therefore, we hypothesized that mitochondrial dysfunction leading to HRI activation might be upstream of ATF4-ISR induction by BTM-3528.

### BTM-3528 induces OMA1-dependent mitochondrial fragmentation

Having established that BTM compounds trigger the ATF4-integrated stress response together with upregulation of genes associated with mitochondrial stress, we next asked whether BTM compounds induce mitochondrial dysfunction. Several recent studies have shown that agents that depolarize mitochondria and block mitochondrial ATP synthesis activate the mitochondrial protease OMA1. Increased OMA1 activity induces the cleavage of OPA1, a dynamin-like protein that controls inner membrane fusion and cristae shape, leading to mitochondrial fragmentation. (25–27). Consequently, we generated OMA1-knockout HCT116 cells using CRISPR editing and performed live imaging of cells stained with Mitotracker Green (MTG) to assess the effects of BTM-3528 on mitochondrial structure in wild-type (WT) and *OMA1^−/−^* cells treated with 3 μM BTM-3528. BTM-3528 induced mitochondrial fragmentation, as indicated by the reduction in both mitochondrial aspect ratio and form factor in WT cells, but not in *OMA1^−/−^* cells (**Fig. 4A and B).**

**Fig. 4.**
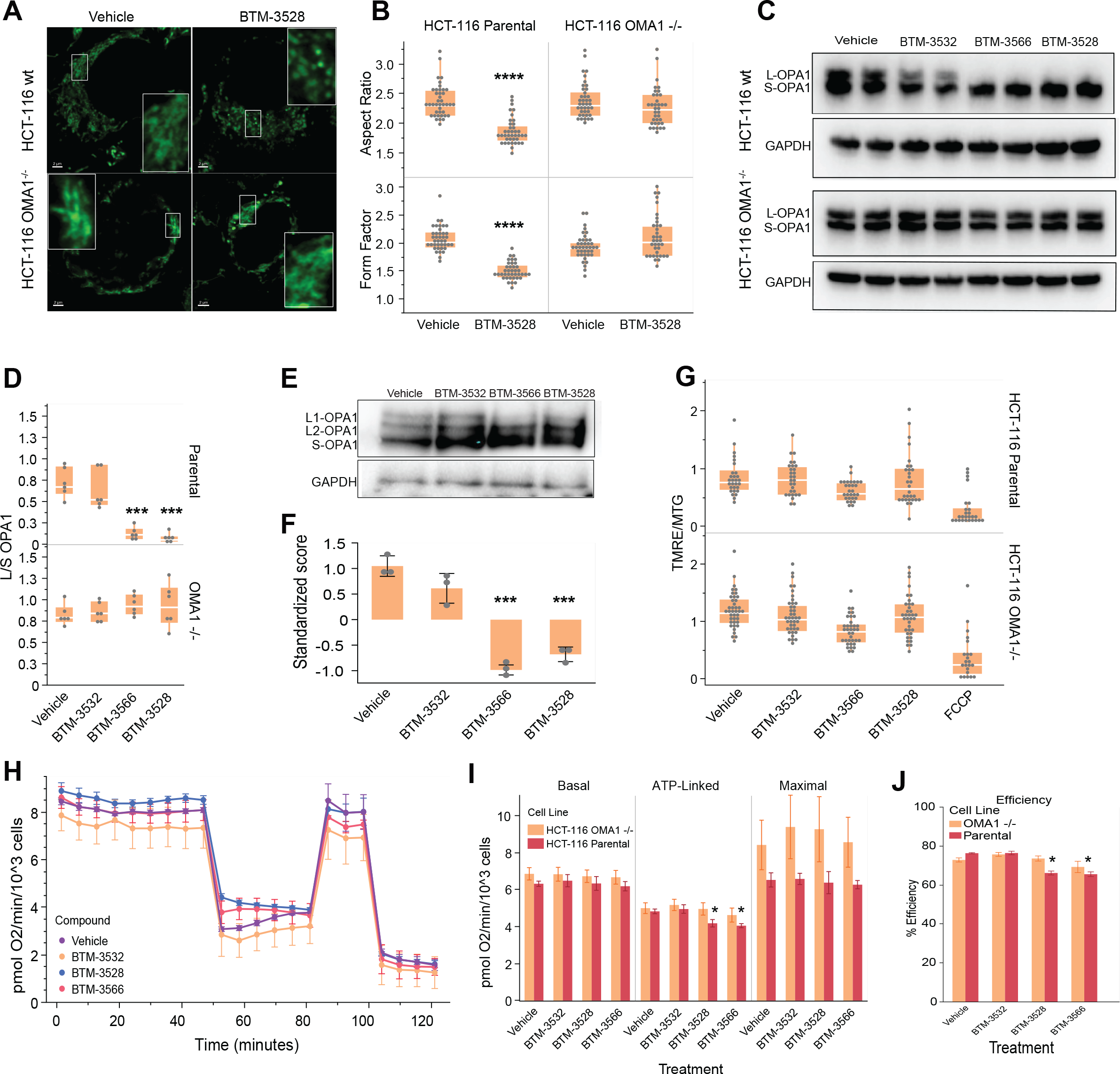
BTM compound treatment fragments the mitochondria and induces OPA1 cleavage in an OMA1 dependent manner. **A)** Representative images from HCT116 cells stained with mitotracker green (MTG) and subjected to live-cell imaging using confocal laser microscopy. **B)** Mitochondrial Aspect Ratio and Form factor are reduced following treatment with BTM-3528. Aspect Ratio and Form Factor are the average of 4 independent experiments, n= 5 cells and 50 – 70 mitochondria identified per cell, t-test, ***, P<.005) **(C, D)** Representative Western blot analysis and quantification of BTM compound induced OPA-1 long forms (L) cleavage to short isoforms (S). HCT-116 Parental cells (+/+) and HCT-116 0MA1 −/− treated for 4 hours with BTM compounds; Bar graph plotted as the average +/− SEM, n=3. ****p<0.0001. **(E, F)** Western blot analysis and quantification of the ratio of the GAPDH normalized L1 OPA-1 to L2 isoforms in parental HCT-116 cells treated with compound for 30 minutes. Bar graph plotted as the Standardized Score calculated for each experiment. ** p<.005 (n=3) **G)** Mitochondrial Membrane Potential in HCT-116 cells treated with 3 μM BTM-3532, BTM-3528 and BTM-3566 for 4 hours then stained with TMRE and MTG. All images were quantified using merged channels indicating colocalization of the TMRE and MTG pixels. FCCP is a used as a positive control. **H)** Respirometry of HCT-116 cells treated with compounds for 30 minutes. All data are plotted as the mean +/− SD (n=3) **I)** Quantitation of the Basal, ATP linked and uncoupled OCR rates and **J)** Bioenergetic efficiency in HCT-116 parental and OMA1 −/− cells. All data are the mean +/− SD, n=3, *, p<.01

Mitochondrial fragmentation can be associated with cleavage of proteins that are required for maintenance of a fused state. OPA1 has 5 isoforms detected by Western blot (bands a-e) and isoform content is sensitive to OMA1 activation.(26–28) OMA1 activation cleaves the long isoforms of OPA1 (L-OPA1) to short OPA1 isoforms (S-OPA1), resulting in mitochondrial fragmentation (22–25). L-OPA1 levels were reduced by more than 90% in parental HCT-116 cells, but not in *OMA1^−/−^* HCT-116 cells following 4-hour incubation with 3 μM BTM-3528 or BTM-3566, but not inactive BTM-3532 (**Fig. 4C and D).** L-OPA1 levels were significantly reduced as early as 30 minutes after treatment with BTM-3528 or BTM-3566 (**Fig. 4E and F**). As noted, apoptosis does not occur upon treatment of HCT-116 cells (**Figure 1F**), indicating that the observed extent of OMA1 activation and OPA1 cleavage are not sufficient to induce apoptosis.

Previous studies have demonstrated that a decrease in mitochondrial ATP production, either achieved by depolarization/uncoupling or by blocking electron transfer (25,26,31), can induce OMA1 activation. Therefore, we determined whether BTM compounds could depolarize mitochondria or directly inhibit electron transfer. WT and *OMA1^−/−^* HCT116 cells were treated with BTM-3528, BTM-3566, or the inactive analog BTM-3532 for 3 hours. Live cells stained with TMRE, a dye reporting on real time changes in mitochondrial membrane potential, and MTG, a dye used to visualize mitochondria mass in a membrane potential independent manner. Co-staining with MTG is used to control for changes in TMRE staining that are not caused by differences in membrane potential (i.e. changes in mitochondrial size or mitochondria moving outside the focal plane). While the mitochondrial uncoupler FCCP strongly reduced the TMRE/MTG ratio, BTM compounds had no effect, consistent with a lack of mitochondrial depolarization (**Fig. 4G**). To further evaluate effects on mitochondrial function, we determined whether acute changes in basal oxygen consumption rate (OCR) occurred after treatment. After 30-minute exposure to BTM-3528 or BTM-3566 in WT cells, no significant changes occurred in basal oxygen consumption in WT or *OMA1^−/−^* cells (**Fig. 4H and I).** ATP-linked respiration and bioenergetic efficiency were mildly decreased and further reduced by 4 hours of treatment in WT but not in *OMA1^−/−^* cells (**Fig. 4I and J; Supplementary Fig. S5A and B).** Thus, effects on respiration from BTM-3528 and BTM-3566 are dependent on the presence of OMA1 but do not result from direct inhibition of electron transport or uncoupling of mitochondrial respiration.

### OMA1-dependent cleavage of DELE1 leads to HRI activation and apoptosis

Having established that BTM compounds induce OMA1 activation without acting as inhibitors of mitochondrial electron transport, we next sought to define the effector pathway leading to lymphoma cell death. In addition to cleavage of OPA1, activation of OMA1 is also known to induce the cleavage of the mitochondrial protein DELE1 and subsequent release of cleaved DELE1 from the mitochondria to the cytosol, where it binds to and activates HRI (32,33). To investigate the role of OMA1, DELE1, and OPA1 as effectors of compound activity in DLBCL, we engineered clonal BJAB cell lines with homozygous deletions of *OMA1* or *DELE1*, or where the OMA1-cleavage site in L-OPA1 was deleted (*OPA1^ΔS1/ΔS1^*). Consistent with our results in HCT116 cells, BTM-3566 failed to induce L-OPA1 cleavage in *OMA1^−/−^* BJAB cells (**Fig 5A**). Processing of L-OPA1 after exposure to BTM-3566 was also blocked in *OPA1^ΔS1/ΔS1^* cells, whereas the deletion of *DELE1* had no effect on the processing of L-OPA1 **(Fig. 5A)**. Activation of the integrated stress response by BTM-3566, as indicated by phosphorylation of eIF2α and induction of ATF4, was lost in OMA1- and DELE1-knockout cells but preserved in *OPA1^ΔS1/ΔS1^* cells unable to undergo OMA1-dependent OPA1 cleavage (**Fig. 5A**). In agreement with these findings, deletion of either *OMA1* or *DELE1* protected BJAB cells from BTM-3566 and BTM-3528 induced apoptosis, whereas *OPA1^ΔS1/ΔS1^* BJAB cells remained fully sensitive (**Fig. 5B, Supplementary Fig. S6A-C**). Thus, activation of OMA1 and cleavage of DELE1 are required for BTM-3566 and BTM-3528-induced activation of the ISR and apoptosis while OMA1-mediated cleavage of OPA1 is dispensable.

**Fig 5.**
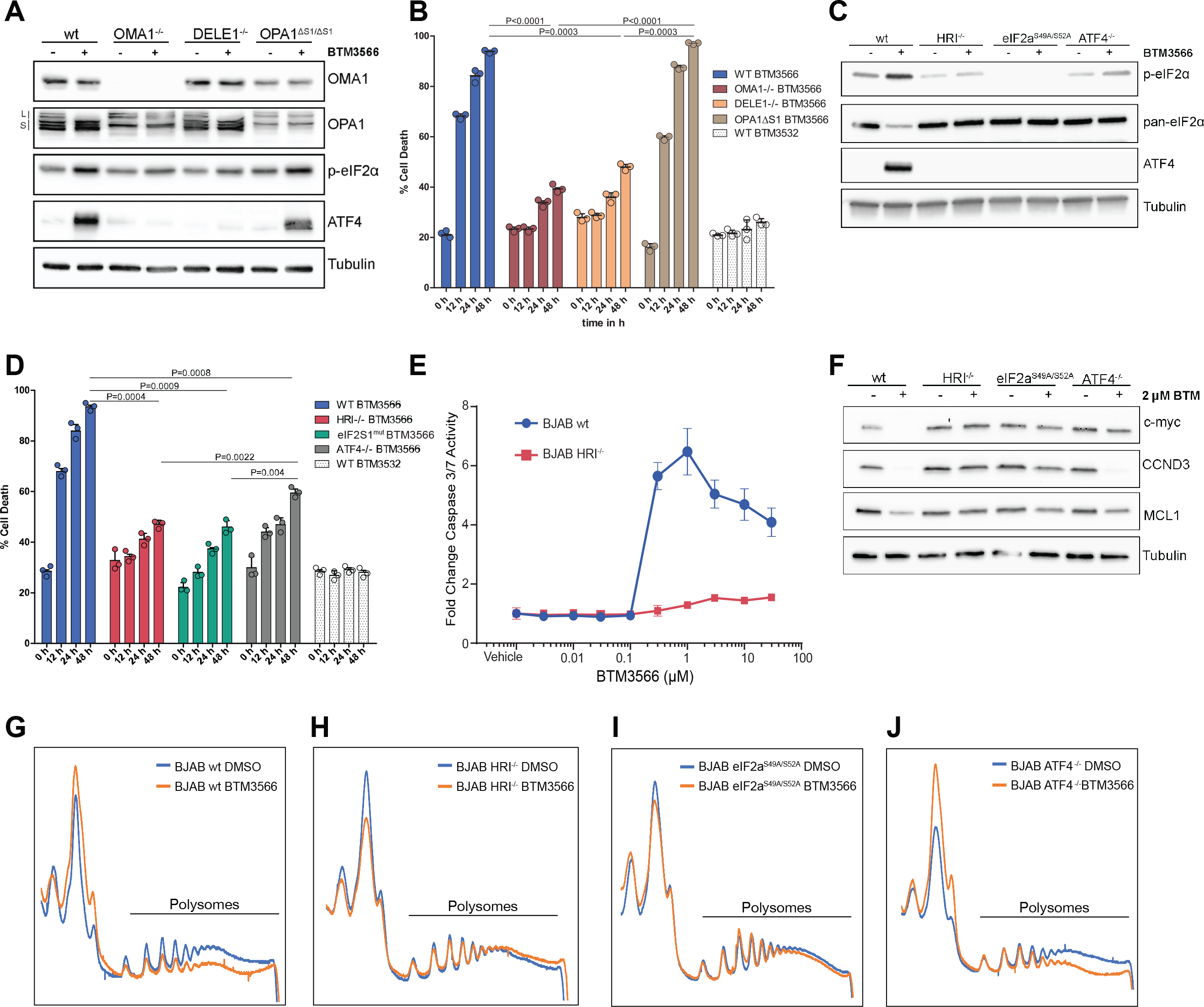
BTM compounds induce the ATF4 ISR via the OMA1-DELE1 mitochondrial quality control pathway. **A)** Model of OMA1 mediated induction of the ATF4 ISR. **B)** Western Blot analysis of OMA1, OPA1, phospho-eIF2α, and ATF4 in parental, OMA1^−/−^, DELE1^−/−^, and OPA1^ΔS1/ΔS1^ BJAB cells following 4 hours of treatment with vehicle or 2 μM BTM-3566. **C)** Cell Death as quantified by Annexin-Propidium Iodide staining in OMA1^−/−^, DELE1^−/−^, and OPA1^DS1/DS1^ BJAB cells following treatment with 2 μM BTM-3566 after 12, 24, and 48 hours of exposure. All data are plotted as the mean +/− SD (n=3). **D)** Western Blot analysis of phospho-eIF2 and ATF4 levels in parental, HRI^−/−^; eIF2α^Ser49/Ser52^ mutant and ATF4^−/−^ BJAB cells following 4 hours of treatment with vehicle or 2 μM BTM-35*66*. **E)** Cell Death as assessed by Annexin-Propidium Iodide staining in parental, HRI^−/−^; eIF2α^Ser49/Ser52^ mutant and ATF4^−/−^ BJAB cells following treatment with 2 μM BTM-35*66* after 12, 24, and 48 hours of exposure. All data are plotted as the mean +/− SD (n=3). **F)** Induction of Caspase 3/7 activity in BJAB wt and HRI^−/−^ cells upon treatment with BTM3566. All data are plotted as the mean (+/− SD, n=3) as compared to vehicle. **G)** Immunblot of c-Myc, CCND3, and Mcl1 in parental, HRI^−/−^; eIF2α^Ser49/Ser52^ mutant and ATF4^−/−^ BJAB cells following 8 hours of treatment with vehicle or 2 μM BTM-35*66*. **H-K)** Polysome profiles of parental **(H)**; HRI^−/−^ **(I)**; eIF2α^Ser49/Ser52^ **(J)** and ATF4^−/−^ **(K)** treated with 2 μM BTM- 3566 or vehicle for 8 hours

### HRI-eIF2α-ATF4 activation leads to apoptosis in DLBCL cells involving loss of oncoproteins regulated by cap-dependent translation

To further determine how the apoptotic signal generated in mitochondria is relayed, we investigated the role of each component of the ATF4-ISR. We created clonal BJAB lines with *HRI* or *ATF4* knockout and a BJAB cell line with a homozygous knock-in of *eIF2α^S49A/S52A^*, in which the serine residues phosphorylated by HRI are mutated to alanine, preventing phosphorylation of eIF2α by HRI. Following treatment with BTM-3528 or BTM-3566 compounds, neither phosphorylation of eIF2α nor increased ATF4 protein was observed in *HRI^−/−^* and *eIF2α^S49A/S52A^* cells. In contrast, phosphorylation of eIF2α was maintained in ATF4-deficient cells (**Fig. 5C, Supplementary Fig 6D**). *HRI^−/−^* and *eIF2α^S49A/S52A^* BJAB cells were protected from compound-induced apoptosis. *ATF4^−/−^* cells were also protected from compound-induced apoptosis, but the effect was somewhat weaker at later timepoints (**Fig. 5D; Supplementary Fig. 6C**). Confirming these data, we found that Caspase 3/7 activation in response to BTM-3566 did not occur in *HRI^−/−^* cells **(Fig. 5E)**.

Known ISR effects downstream of eIF2α phosphorylation include downregulation of 5’-cap-dependent mRNA translation with concomitant decreases in the levels of short lived oncogenic proteins and increased μORF-mediated translation of ATF4 and other stress effectors.(17,34,35) We investigated the impact of treatment on the abundance of short-lived oncogenic proteins essential for DLBCL cells and found that BTM-3528 treatment depleted c-MYC and Cyclin D3 (CCND3) proteins and reduced the expression of the anti-apoptotic protein MCL1 after 8 hrs of treatment (**Fig 5F**; **Supplementary Fig. S6 B,C**). To quantify cellular protein synthesis, we performed polysome gradients, which revealed that upon treatment with BTM-3566, polysomes from WT and *ATF4^−/−^* BJAB cells exhibited a shift towards smaller size, indicating global attenuation of cap-dependent mRNA-translation **(Fig. 5G and J)**. In contrast, cells with knock-out of *HRI* or knock-in of *eIF2α^S49A/S52A^* had preserved polysome size (**Fig. 5H and I**). In *HRI^−/−^* or *eIF2α^S49A/S52A^* BJAB cells, the effect of BTM-3566 on c-MYC, CCND3, and MCL-1 was fully rescued, whereas in *ATF4^−/−^* cells the expression of c-MYC and MCL-1 was largely preserved, but CCND3 was still lost (**Fig 5F**).

### The mitochondrial protein FAM210B suppresses BTM-3528 and BTM-3566 activity

We sought to understand factors that modulate the activity of BTM-3528 and BTM-3566. Therefore, we extended our screen for compound activity to a larger panel of cell lines (**Supplementary Fig. S7**). Each cell line was treated in duplicate with BTM-3528 to establish a dose-response. The activity area (Area Under Curve, AUC) for each dose-response was then computed (**Supplementary Table S3**). The AUC response data were analyzed for correlation to genomic alterations or gene expression using data from the Cancer Cell Line Encyclopedia ((36)). No single mutations were found to be associated with compound activity in 284 cell lines for which both gene expression and genomic data were available.

We noted a striking relationship between expression of the mitochondrial protein FAM210B and response to BTM-3528 across all cell lines tested (**Fig. 6A, Supplementary Tables S4 and S5, Supplementary Figure S8**). *FAM210B* expression levels were significantly lower in drug-sensitive cell lines (**Fig. 6B**). Notably, the most responsive cell lines were derived from B-cell lymphomas (DLBCL, Burkitt’s and Mantle cell lymphoma) which also had the lowest levels of *FAM210B* expression (**Fig. 6B and C**). An analysis using the BioGPS database revealed that FAM210B expression levels in normal human blood cells are low compared to solid tumor cells and that expression is lowest or absent in centrocytes, centroblasts and naïve B-cells (**Fig. 6D**) (37,38). This suggests that low FAM210B levels found in DLBCL reflect the cell of origin and not mutation or derangement in gene expression.

**Fig. 6.**
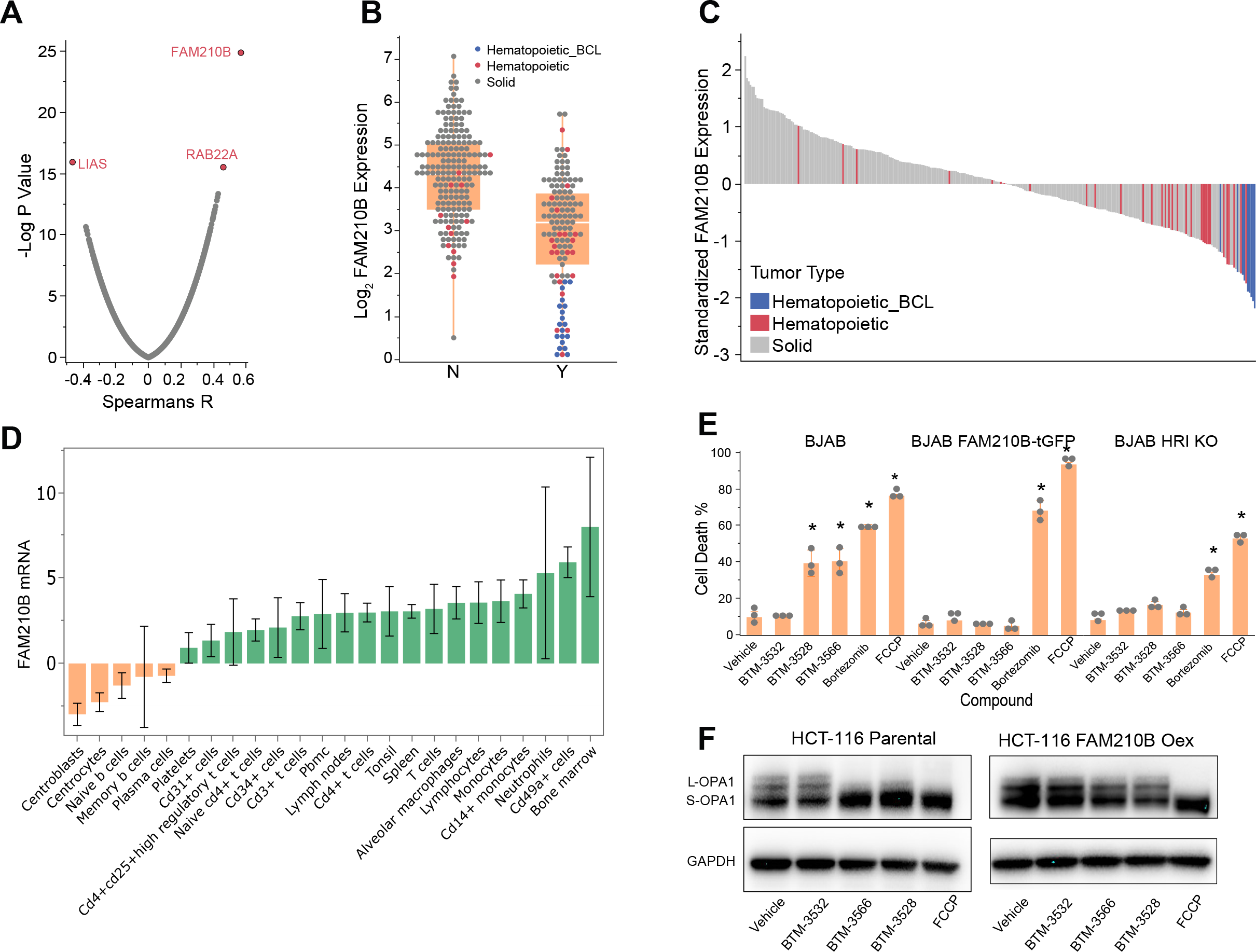
*FAM210B* mRNA expression is correlated with and regulates cellular response to BTM compounds. **A)** BTM-3528 activity was assessed in 406 solid and hematopoietic tumor cell lines. Gene expression was correlated to AUC across 284 cell lines that found with gene expression data in the CCLE database. The data are expressed as the Spearman’s correlation coefficient vs the -LogP value for each gene. **B)** Log2 expression of FAM210B in cell lines denoted as responsive or not to BTM-3528 (n=284). Cells were classified as responder (AUC > 3.25) or non-responder (AUC < 3.25) *** p<.001. • Solid tumor lines; • Hematopoietic tumor lines; • B-cell lymphoma (BCL) Lines. **C)** Waterfall plot of *FAM210B* mRNA expression data for screened cell lines. Data are plotted by descending standardized *FAM210B* expression level. **D)** FAM210B gene expression levels in normal human blood cells. All data from BioGPS (http://ds.biogps.org/?dataset=BDS_00001&gene=116151) **E)** Parental, FAM210B-tGFP or *HRI^−/−^* BJAB cells were tested for sensitivity to BTM-3528, BTM-3566, BTM-3532, Bortezomib and FCCP. Cell death was determined by Annexin and YOYO staining. **F)** OPA1 cleavage was determined in HCT-116 parental and FAM210B-tGFP cells following a 3-hour treatment with 3 μM BTM compound**s**

To determine whether FAM210B suppresses the effects of BTM-3528 and BTM-3566, we engineered BJAB cells to stably overexpress FAM210B-tGFP and compared them to wt and *HRI^−/−^* cells. Bortezomib and FCCP were included as controls to assess the specificity of FAM210B on preserving cell viability when ATF4-ISR is induced via alternative methods of activation. Strikingly, FAM210B-tGFP expression completely suppressed the activity of BTM-3528 and BTM-3566 but had no effect on bortezomib or FCCP-induced cell death **(Fig. 6E).** Importantly, knock-out of *HRI* reduced the activity of both bortezomib and FCCP, confirming the common utilization of HRI in induction of the ATF4 ISR and subsequent apoptosis.

To determine the effects of FAM210B on OMA1 activation, we compared the ability of BTM-3528, 3566, and the inactive control BTM-3532 to activate OMA1 in the presence or absence of FAM210B-tGFP overexpression. As expected, L-OPA1 cleavage in WT HCT-116 cells was observed in the presence of BTM-3528 or BTM-3566 but not BTM-3532 (**Fig. 6F**). L-OPA1 cleavage did not occur in HCT-116 cells stably expressing FAM210B-tGFP (**Fig. 6F**), indicating that FAM210B acts upstream of OMA1 to suppress the effect of compounds on OPA1 cleavage. FCCP-induced cleavage of OPA1 was unaffected by FAM210B-tGFP. We also compared the effects of FAM210B expression on ATF4 protein expression using BTM-3566 and alternative inducers of the ATF4 ISR. FAM210B expression robustly suppressed ATF4 induction by BTM-3566 but not suppressed by amino acid starvation, tunicamycin, bortezomib, or ONC201 treatments **(Supplementary Fig. S9)**, suggesting a unique mechanism. In addition, compound-dependent decreases in basal, ATP-linked, and maximal mitochondrial respiration were blocked in cells overexpressing FAM210B-tGFP (**Supplementary Fig. S10A and B**).

## DISCUSSION

Tumors acquire the ability to overcome normal growth control mechanisms (*36*). Mitochondria play a crucial role in this process as central hubs controlling cell survival in response to proapoptotic stimuli that activate the intrinsic apoptotic pathway (*37*). The novel pharmacology described here presents a therapeutic opportunity. Importantly, therapeutic activity of BTM-compounds appears to span a diversity of genetic backgrounds in DLBCL and other B-cell lymphomas that differ in cell-of-origin, presence of *MYC* rearrangements and other genetic alterations. Collectively, our data suggest a model in which BTM compounds activate the mitochondrial protease OMA1 only in the absence of FAM210B. Activated OMA1 cleaves DELE1, whose processed form relays the signal to the cytosol and the HRI-eIF2α-ATF4 effector pathway (**Fig. 7**). In DLBCL tumors, this sequence of events leads to robust and rapid apoptosis and tumor regression. The biological reason for robust sensitivity of DLBCL lines is unclear but could be related to low expression of FAM210B, which could also serve as a biomarker for the selection of patients most likely to respond to OMA1 activators.

**Figure 7:**
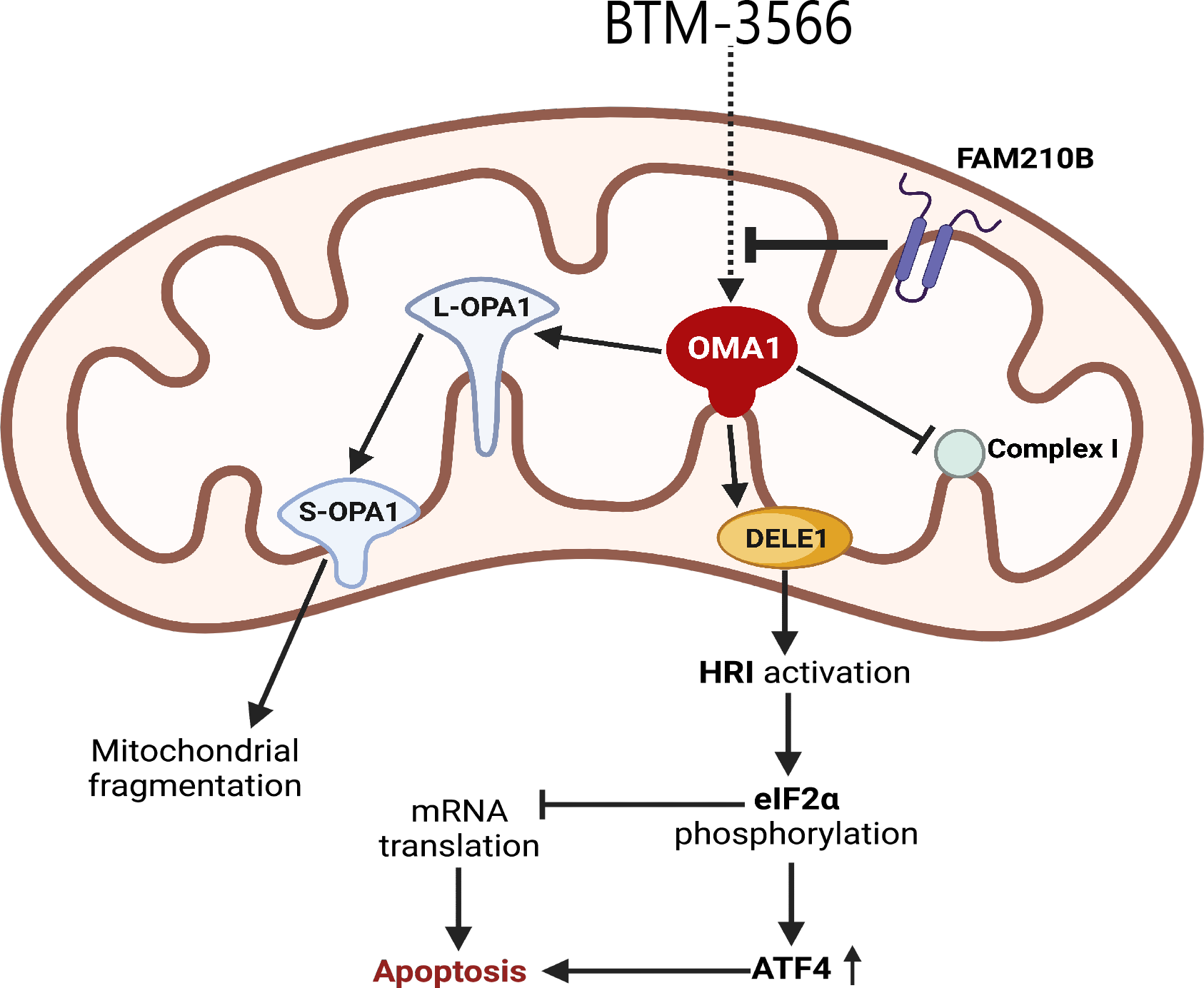
Model for the mechanism of action of BTM compounds. BTM compounds induce the activation of OMA1 leading to the cleavage of DELE1 and OPA1. OPA1 activation leads to fragmentation of the mitochondrial network. DELE1 cleavage leads to activation of HRI and the downstream effector pathways leading to cell death in DLBCL cell lines.

The downstream consequences of OMA1 activation are two-fold: cleavage of DELE1 followed by activation of the ATF4 ISR and fragmentation of the mitochondrial network associated with cleavage of OPA1. The therapeutic effects of BTM compounds in DLBCL are linked to activation of the ATF4 ISR, resulting in inhibition of global cap-dependent translation and increased ATF4-dependent transcription. It is notable that BTM-compounds are uniformly effective in DLBCL lines that have differing sensitivities to venetoclax or variable expression of components of the intrinsic apoptotic pathway, including MCL1, BCL2 and BCL-XL (39–41). The exact molecular context that makes DLBCL uniformly sensitive to induction of the ATF4 ISR by BTM-compounds are not fully understood. Persistent eIF2α phosphorylation can sensitize cells to apoptotic stimuli by reducing the levels of anti-apoptotic proteins such as MCL1, BCL2 and XIAP (10–12,15) as well as upregulating expression of proapoptotic genes such as TRB3 (16) and BIM (14). Translation inhibition is also responsible for induction of ATF4-regulated gene transcription. Drug regulated ATF4 target gene such as DDIT3 (CHOP) are known to promote cell death through a variety of mechanisms in the context of the ER-stress response (42–44). Thus, additional results are needed to identify factors distinguishing BTM-compound sensitive from resistant tumors and whether particular agents capable of inducing apoptosis (e.g. the BCL2 inhibitor venetoclax) are likely to synergize with BTM-compounds in specific contexts.

Cleavage of OPA1 and fragmentation of the mitochondrial network are prominent outcomes of BTM-compound activity. OPA1, a dynamin-like like protein involved with fusion of the inner mitochondrial membrane, is intimately involved in maintenance of cristae structure and mitochondrial morphology (27,29,30). Controlled OMA1-dependent proteolysis of L-OPA1 to S-OPA1 has been implicated as an early event associated with apoptosis (26,45–48). Our findings suggest that BTM-compound activation of OMA1-dependent OPA1 cleavage is neither necessary nor sufficient to activate apoptosis. This is particularly true in solid tumor lines where cell growth arrest, but not apoptosis, is the outcome. This suggests that BTM-compound induced cleavage of OPA1 and fragmentation of the mitochondrial network are not universally capable of inducing tumor regression, at least in the contexts tested here.

The precise mechanism by which BTM-compounds induce OMA1 activation remains unclear. The compounds had no effect on mitochondrial electron transport or membrane potential suggesting an alternative mechanism for activation of OMA1. That mitochondria are central to the pharmacologic effects of BTM-compounds is reinforced by the inverse correlation between cell line sensitivity and expression of the mitochondrial protein FAM210B. Overexpression of FAM210B in both BJAB and HCT116 cells conferred resistance to BTM-compound induced activation of the ATF4 ISR while having no effect on other inducers of the ISR (tunicamycin, nutrient deprivation and bortezomib) or compounds that affect mitochondrial membrane potential (the uncoupler FCCP) or mitochondrial proteostasis (the CLPP1 activator ONC201). As such, there must be multiple pathways inside mitochondria that, when perturbed, lead to activation of the ATF4 ISR but only a FAM210B-regulatable process is affected by BTM-compounds.

The physiological role of FAM210B appears to depend on cellular context. FAM210B is involved in high-capacity heme biosynthesis occurring during erythropoiesis, acting to increase the importation of iron into the mitochondria. FAM210B is not required for basal heme biosynthesis (49–51). FAM210B may have broader activity controlling cellular metabolism unrelated to heme biosynthesis but tied to its ability to act as a scaffold protein for other mitochondrial membrane or matrix proteins. Reduction in FAM210B expression using siRNA leads to increased mitochondrial respiratory capacity, decreased glycolysis and an aggressive metastatic tumor phenotype (52). FAM210B has also been described as a tumor suppressor, with lower FAM210B levels associated with poor prognosis in patients with renal, cervical and lung cancers (52,53).

Multiple therapeutic strategies have been utilized to induce eIF2α phosphorylation and the ATF4 ISR as treatment for cancer. The 26S proteasome inhibitor bortezomib and the novel CLPP1 activator ONC201 rely on activation of HRI to phosphorylate eIF2α, with subsequent induction of ATF4 and CHOP (54–63). Our data provide further support for the induction of ATF4 ISR, in this case through pharmacologic activation of OMA1 as an orthogonal strategy for inducing cancer growth arrest and apoptosis.

## MATERIALS AND METHODS

### Cell line compound testing

Screening of tumor cell lines was performed by Crown Bioscience (OmniScreen Platform). The protocol estimated the growth of each cell line from the initial plating density. Cells were plated at a starting density of 4×10^3^ cells/well and incubated for 24 hours. BTM-3528 was prepared as a 10×solution of test article with a final working concentration of 30 μM of test article in media with nine 3.16-fold serial dilutions, following addition of BTM-3528, the plates were incubated for 96 hours at 37°C with 5% CO2. Final cell numbers were determined using the Cell-Titre Glo assay (Promega). The absolute IC_50_ (EC_50_) curve was fitted using nonlinear regression model with a sigmoidal dose response.

**Caspase 3/7 activity** was determined following treatment of cells with BTM-3528 or BTM-3566. using the Caspase-Glo® 3/7 assay (Promega).

### Annexin V Apoptosis Assay

To quantify apoptosis, BJAB cells were cultivated in a 96-well format and treated with BTM compounds for the indicated time intervals. Cells were washed twice with ice cold PBS and resuspended in 1x Binding Buffer (BD Biosciences) at a concentration of 1 × 10^6^ cells/ml. To 0.5 × 10^5^ cells 2.5 μl of Annexin V-APC or Annexin V-FITC (BD Bioscience) were added and incubated for 15 min at RT in the dark. Cells were washed once with Binding Buffer and the pellets were resuspended in 100 μl Binding Buffer containing either 2 μl PI (50 μg/ml) or 2 μl DAPI (1 mg/ml). Cells were analyzed on a Cytoflex S (Beckman Coulter).

### Transcriptomic Profiling

The human colon adenocarcinoma cell line HCT116 was used to evaluate the effects of BTM compounds on gene expression. To fully evaluate the effects of the compound on cell-cycle controlled genes, cells were synchronized prior to BTM compound treatment. HCT-116 cells were first blocked in S phase by treatment with thymidine. After 24 hours, the thymidine containing media was removed and replaced with media containing nocodazole to block cells in M phase in a high degree of synchrony. Cells were then released into G1 either in complete medium or complete medium plus 10, 1, or 0.1 μM BTM-3528. Cells were harvested at five time points: 1, 2, 4, 6, and 8 hours after release into G1 phase, and mRNA extracted for Illumina RNA-seq. Three replicates of each concentration and time point along with time point specific controls (i.e., cells without compound) were collected for RNA-sequencing.

### Xenograft models

Human cell line xenograft models were established using SU-DHL-10 cells (ATCC Cat# CRL-2963). Cells were grown in RPMI 1640 supplemented with 10mM HEPES buffer, /L-Glutamine, 1mM Na pyruvate, 4500 mg/L Glucose, 15% Non-Heat-Inactivated Fetal Bovine Serum and Penicillin/Streptomycin. Cells were harvested by centrifugation and resuspended in cold 50% serum-free medium: 50% Matrigel to generate a final concentration of 2.50E+07 trypan-excluding cells/mL. Female Envigo SCID beige mice (C.B-17/IcrHsd Prkdc^scid^lyst^bg-j^) were implanted subcutaneously high in the right axilla on Day 0 with 5×10^6^ cells/mouse. Mice were randomized into study groups based on tumor volume with a mean tumor burden for each group of 150mm^3^. BTM-3566 was prepared as a solution in dosing vehicle containing 5% NMP, 15% PEG400, 10% Solutol, and 70% D5W. All mice were dosed once daily by oral gavage for 21 days. The final dose concentration was 4 mg/ml, and the dose volume was 5 mL /gram. Tumor volume and body weights were determined every 3^rd^ day. All mice were dosed according to individual body weight on the day of treatment.

For the patient derived xenograft models, all tumors were sourced from Crown Bio. Models were established in female mice with an average body weight of 25 grams. Balb/c nude from GemPharmatech Co.,Ltd (LY0257); NOD SCID mice from Shanghai Lingchang Biotechnology Co., Ltd (Shanghai, China) (LY2214, LY2264, LY2345, LY3604, LY6701) or NPG/NOD/SCID from Beijing Vital Star Biotechnology Co,Ltd (LY6933, LY6934) mice. Each mouse was inoculated subcutaneously in the right flank region with fresh tumor derived from mice bearing established primary human cancer tissue. Mice were randomized into a vehicle or treatment group with a mean tumor burden of 200 mm^3^. All mice were dosed once daily by oral gavage for 21 days. Tumor volume and body weights were determined three times per week.

All animal studies were conducted under strict ethical and animal welfare guidelines following an approved IACUC protocol from the host institutions. Although the studies were not conducted in accordance with the FDA Good Laboratory Practice regulations, 21 CFR Part 58, all experimental data management and reporting procedures were in strict accordance with applicable industry practices Guidelines and Standard Operating Procedures.

#### Confocal live cell imaging

HCT-116 cells were plated in a 35 mm dish containing a 14 mm, uncoated coverslip (MatTek). The next day, cells were stained with McCoy media containing 200nM Mitotracker green (MTG) and 15nM of tetramethyl rhodamine, ethyl ester (TMRE) for 40 minutes. Cells were then treated with 3μM BTM-3528 or 4μM FCCP in McCoy containing 15nM TMRE, but lacking MTG, and incubated for 3h. Imaging was perfumed using a 63x objective and in a Zeiss LSM880 confocal microscope with Airyscan mode.

#### Mitochondrial membrane potential

Cells were plated in black clear-bottom 96 wells plate and stained with McCoy media containing 200nM MTG and 15nM of TMRE for 40 minutes. Cells were then treated with 3μM BTM-3528 or 4μM FCCP in McCoy containing 15nM TMRE, but lacking MTG, and incubated for 30 minutes. MTG and TMRE signal was obtained by imaging in two wavelengths (MTG – Excitation 460-490 nm and Emission 500-550 nm; TMRE – Excitation 560-580 nm and Emission 585-605 nm) using a PerkinElmer Operetta microscope system with 40x lens. Five fields of each well were analyzed and TMRE/MTG fluorescence ratio was determined by image processing with Harmony software.

#### Image analysis

Mitochondrial morphology was assessed using Fiji/ImageJ software and a Trainable Weka Segmentation plugin. Mitochondrial membrane potential, TMRE/MTG fluorescence ratio was calculated from segmented mitochondrial structures obtained by MTG channel. One-way ANOVA and Tukey’s multiple comparison test were used for statistical analysis; P-values ≤ 0.05 (*) were considered significantly different.

#### Mitochondrial Respirometry

Respirometry assays were run on a Seahorse Extracellular Flux Analyzer (Agilent). HCT116 cells were seeded at 14,000 cells/well using XF96 well microplates and incubated overnight (37°C and 5% CO2) in McCoy’s 5a Modified Medium culture medium with 10% FBS. Before the respirometry assay, cells were washed with assay medium: DMEM with 10mM glucose, 2mM glutamine, 1mM pyruvate, 5mM HEPES, and 10% FBS (pH 7.4). BTM compounds were tested at a final concentration of 3μM, and cells were either acutely treated during the assay or pretreated for 4 hours before the assay. In pretreatment experiments, compounds were added in complete medium and incubated at 37°C and 5% CO2. Compounds injected during the assay included 2μM oligomycin, 1μM FCCP, and 2μM of antimycin A and rotenone. Upon completion of each respirometry assay, the cells were stained with 1μg/mL Hoechst and cell number was measured with an Operetta High-Content Imaging System. The respirometry well level data (pmol O2/min) was normalized to cell number per well (pmol O2/min/10^3^ cells) in each assay.

## Supporting information

RNA Seq Data

## SUPPLEMENTARY MATERIALS

### Materials and Methods

**Table S1**: Cell Line Screening

**Table S2**: DLBCL Cell Line Screening

**Table S3**: Crown Screening Data

**Table S4**: Common genes predictive of response to compound.

**Table S5**: Biomarker Screening data analysis Spearmans Analysis

**Table S6**: RNASeq_BTM3528_Data and Analysis

## ACKNOWLEDGEMENTS

The authors would like to thank Dr. Josh Rabinowitz of Princeton University for helpful scientific discussions, comments, review and editing of this manuscript. Dr. Alexander van der Bliek, at the UCLA David Geffen School of Medicine UCLA for the HCT-116 OMA 1−/− cell line.

## AUTHOR CONTRIBUTIONS

Conceptualization: MK, MLR, AdS, MH

Methodology: MK, MLR, MO, AdS, MJK, SS, MH, AD, AK, MG, AS

Investigation: MK, MLR, MO, AdS, MJK, SS

Visualization: MK, MLR, MO, AdS

Supervision: MK

Writing – original draft: MK, AdS, MH, MLR, DW

## DATA AND MATERIALS AVAILABILITY

RNASeq data have been deposited in NCBI’s Gene Expression Omnibus. A reviewer access link has been generated and can be obtained from the authors. The R-Code used to perform the bioinformatic analysis is available upon request from the authors. All other data are available in the main text or the supplementary materials. BTM3528 and BTM3566 are available upon request through a materials transfer agreement (MTAs) with Bantam Pharmaceutical.

## Supplementary Materials for

### Supplemental Methods

#### Cell lines

BJAB cells were obtained from the German Collection of Microorganisms and Cell lines (DSMZ #ACC 757). HCT-116 cells were purchased from the American Type Culture Collection (CCL-247).

#### Genomic engineering of BJAB cells

BJAB cells were obtained from the German Collection of Microorganisms and Cell lines (DSMZ #ACC 757). Alt-R crRNA, Alt-R tracrRNA and Alt-R Cas9 Nuclease V3 were purchased from IDT. The genomic crRNA target sequences were: ATF4: 5’-GCGGGCTCCTCCGAATGGC-3’, HRI: 5’-ATAGTCGAGAGAAACAAGCG-3’, eIF2S1: 5’-TTCTTAGTGAATTATCCAGA-3’, OMA1: 5’-CTGGAAGTAAGTCCAATCAC-3’, OPA1: 5’-GCGTTTAGAGCAACAGATCG-3’, DELE1(knock-in):5’-GAAAGGAGTGTTGTAAGACT-3’, DELE1 (knock-out): 5’-ACTGGGACCTAGCCTCTGGA-3’. For HDR-mediated knock-ins IDT ultramers with the following sequences were used as DNA donors. The base substitutions introduced for mutation of S49 and S52 are in **bold capital**: 5’-a*c*ttacctttttctttgtccaccctaatgacaaccacacactcattcctgccaattcggatgagtttgttgatagaacggatacgtcttct**TG C**taattc**TGC**aagaagaatcatgccttcaatg*t*t-3’. For the knock-in of a HA-tag into the DELE1 locus the following template was used with the HA-tag and linker in uppercase: 5’-t*g*gaaagggttgctgatgctctatttttgttatccaacctcctgcccgctctggcccctaagaggcaccagggactatgttttatctcaccT TACCCAGCGTAGTCTGGGACGTCGTATGGGTATGAGCCGCCTCCGCTCCCTCCGCCgccaaaacctaaccttacaacactcctttccagtgggtaggggtgtg*g*g-3’. For the specific deletion of the S1 cleavage site of OPA1 the following template was used (the 10 amino acids deletion around the S1 cleavage site is marked in **bold**): c*a*aagaagtaaaaactttaaaaaatctttcaagactacctacatgaacaattctcttttacacttacctttctaaaatgcttgtcacttt**ct**tccg gagaaccttaaagataaatatgcaaccct*t*t (* marks phosphorothioate bonds). Cas9 RNPs were made by incubating 105 pm Cas9 protein with 120 pmol Alt-R crRNA:tracrRNA duplex in 5 μl volume at 25 °C for 10 min immediately prior to electroporation. 2.5*10^6^ BJAB cells were electroporated using the Lonza Nucleofector 4D and 3.85 μM IDT electroporation enhancer. For HDR-mediated knock-in of the eIF2 S49A S52A variant, 3.7 μM single-stranded oligo DNA nucleotide (ssODN) was added to the electroporation mix. Single clones were obtained by single-cell sorting of electroporated BJAB into 96-well plates and validated by TIDE or ICE (*70*) analysis of single clones to identify homozygous clones.

#### Cell Cycle Analysis

HCT-116 cells were plated into black-walled poly-L-lysine coated microclear tissue culture plates and incubated overnight. Cells were treated with compound dissolved in DMSO and diluted into McCoys modified media with 10% fetal bovine serum. The cells were incubated with compound for 24 hours after which they were treated with EdU to label newly synthesized DNA and allow for positive identification of S-phase cells. The cells were washed then fixed with 4% formaldehyde and permeabilized with 0.1% Triton X-100. Following fixation and permeabilization, cells were washed and treated with Click-IT according to the manufacturer’s instructions (Thermo Fisher #C10637). Following Click-IT labeling, the cells are washed and blocked with 3% bovine serum albumin in Dulbecco’s PBS – Ca – Mg. A phosphoHistone H3 antibody (Santa Cruz; cat#sc-8656-R) is diluted 1:50 in blocking buffer and then added to the cells for one hour. The cells are washed and subsequently treated with a solution of blocking buffer containing 1 ug/mL DAPI and goat anti-rabbit antibody (1:200 dilution of Alexa Fluor 647 goat anti-rabbit IgG; Molecular Probes; cat#A21245.). The labeled and stained cells were imaged using an ArrayScan XTi High Content Imager with a 20x/0.4NA Plan-Neofluar objective using three fluorescent channels. Image analysis was performed using Cellomics HCS Studio software. Cell cycle classification and statistical analysis was performed using Knime. The output of the image analysis is: total number of cells per well, total DNA content, specific identification of cells in S-Phase (EdU staining) and identification of those cells at the G2/M boundary (phosphoH3 positive). All datapoints shown are means +/− SD of three biological replicates.

#### Image analysis of ATF4 in HCT-116 cells

HCT-116 are cultured in McCoys 5A modified media with L-Glutamine, 10% FBS and Pen-Strep. Cells were grown to near confluency in a T-75 flask, then harvested using 0.25% Trypsin-EDTA. Harvested cells were washed with Dulbecco’s PBS -Ca -Mg, the resuspended in complete medium to a final concentration of 10,000 cells/mL then dispensed into Cell Carrier Ultra 96 well plates coated with poly-L-Lysine. The plated cells were returned to a 37°C, 5% CO_2_ incubator for 24 hours. For treatment of cells with compound, a solution of 15 mM BTM-3528 was first prepared. A solution of 30 mM BTM-3528 was prepared in 1 mL of McCoys complete media containing 0.2% DMSO then serially diluted in half-log increments in the same media. The plated cells were then removed from the incubator, the media aspirated and replaced with the drug containing treatment media. The cells were incubated for 4 hrs, then fixed with 4% formaldehyde and permeabilized with cold methanol. The cells were washed, and the plates blocked with 5% normal goat serum in Dulbecco’s PBS +Ca+Mg (DPBS+CM). Following blocking the cells were washed in DPBS+CM then incubated for 1 hour with 50 mL/well of a 1:200 dilution of rabbit anti-ATF4 antibody (Cell Signaling technology # 11815). The cells were then washed with DPBS-CM and incubated in 50 mL DPBS-CM containing 5% normal goat serum, 2 mG /mL goat anti-rabbit Alexa Fluor 647 (ThermoFisher #A21245) and 1% DAPI. The cells were washed in DPBS+CM and imaged using a Opera Phenix on confocal with a 63X water immersion objective. The DAPI (405nM) and Alexa (647 nM) channels were then imaged with a minimum of 100 cells per well captured. Image analysis is performed with Harmony v4.5 software (PerkinElmer). Advanced flatfield correction is employed. Nuclei are segmented from DAPI channel images and mean nuclear pixel intensity of the Alexa Fluor 647 ATF4 channel is calculated. For each well, the mean of this value is determined and exported for analysis.

#### OPA1 Western Blotting in HCT-116 cells

HCT-116 cells were harvested, washed three times in PBS then lysed in RIPA buffer [50 mM Tris-HCl, pH 7.5, 0.1 % (w/v) Triton X-100, 1 mM EDTA, 1mM EGTA, 50 mM NaF, 10 mM sodium β-glycerophosphate, 5 mM sodium pyrophosphate, 1 mM vanadate) with protease inhibitor mixture (MilliporeSigma) and phosphatase inhibitors (Roche). Lysates were shaken for 15 minutes at 4 °C followed by centrifugation (12000 xg, 10 minutes at 4°C). Supernatant was collected and protein content was determined by BCA Protein Assay. Cell lysates were diluted in Laemmli sample buffer (100 mM Tris–HCl, 2% SDS, 10% glycerol, 5% β-mercaptoethanol and 0.1% bromophenol blue) and heated at 95 °C for 15 minutes. Proteins were run on 3%-8% SDS-PAGE and transferred onto a PVDF membrane. Membranes were blocked with 5% non-fat milk and detection of specific proteins determined by over-night incubation of membranes with primary antibodies against OPA1 (BD Biosciences; 1:1000), GAPDH (Cell Signaling) or TOMM20 (Abcam), washed then incubated with peroxidase-linked anti-mouse (for OPA1) or anti-rabbit (for GAPDH and TOMM20) secondary antibody for 1h (Calbiochem; 1:10000). Membranes were washed and exposed to a chemiluminescent protein detection system (ChemiDoc, MP Imaging System; Bio-Rad).

#### Western Blotting and ProteomeProfiler™ Arrays in BJAB cells

BJAB cells were harvested and washed in ice cold PBS then lysed in RIPA buffer (Cell Signaling Technology) with complete protease inhibitors cocktail (Roche) and Phostop phosphatase inhibitor mixture (Roche). Lysates were shaken for 15 minutes at 4 °C, sonicated (3 × 10 s) and followed by centrifugation (12000 × g, 10 minutes at 4°C). Supernatant protein content determined by BCA Protein Assay. Cell lysates were diluted in blue loading buffer (Cell Signaling Technology), heated at 95 °C for 5 minutes the applied to 8.5% SDS-PAGE or 4-20% Mini-PROTEAN TGX Stain-Free Precast Gel (Biorad) and transferred onto a Nitrocellulose membrane (Biorad). Membranes were blocked with 5% non-fat milk for 1h and the detection of specific proteins was determined by over-night incubation of membranes with primary antibodies against ATF4 (Cell Signaling Technology; 1:1000), eIF2α (Cell Signaling Technology; 1:1000), p-eIF2α (Cell Signaling Technology; 1:1000), OMA1 (Cell Signaling Technology; 1:1000), OPA1 (Cell Signaling Technology; 1:1000) or α-Tubulin (Sigma, 1:5000). After washing membranes were incubated with anti-rabbit secondary antibody for 1h (Cell Signaling Technology; 1:2000). Membranes were washed and exposed to a chemiluminescent protein detection system (SuperSignal West Pico PLUS, Thermo Scientific). To examine the expression of proteins linked to apoptosis the Proteome Profiler Human Apoptosis Array Kit (R&D Systems, ARY009) was used according to manufacturer’s instructions.

#### Polysome Profiling

All steps were carried out in the cold. 1 × 10^7^ cells were washed in ice-cold PBS containing 0.1 mg/ml cycloheximide, pelleted and resuspended in extraction buffer (20 mM Tris-HCl, pH 8.0, 140 mM KCl, 0.5 mM DTT, 5 mM MgCl_2_, 0.5 % nonidet-P40, 0.1 mg/ml cycloheximide, 10000 IU/ml dalteparin sodium) and incubated for 10 min on ice. After centrifugation for 10 min at 12000 × g, 0.4 ml supernatant was layered onto a 12 ml linear sucrose gradient (10 – 50% sucrose [w/v] in extraction buffer without nonidet-P40) and centrifuged at 4 °C in an SW40Ti rotor (Beckman) at 35000 rpm without brake for 120 min. The gradients were collected, absorbance was measured at 254 nm and profiles were recorded using an optical UA-6 detector (ISCO). For the normalization of the polysome profiles in the x-direction, the maximum of each 80S peak was determined and a common x-start value was set for all 80S maxima. This was followed by y-normalization to enable comparability of individual curves. For this, the area under the curve (AUC) was determined for each run by adding up all OD values. A scaling factor was calculated by dividing each individual AUC to the lowest AUC of the runs within one experiment and all polysome profile data points were divided by the scaling factor.

##### RNAseq Data Analysis

FASTQ files contain the read pairs for a sample along with their base pair quality score. Quality scores are based on the PHRED33 scoring scheme. Illumina RNA-seq read files (FASTQ files) were assessed with FASTQC for read quality (www.bioinformatics.babraham.ac.uk/projects/fastqc/). Quality scores were used to trim the leading and trailing base pairs to retain only those with a minimum PHRED quality score of 30 or better, and with a minimum retained read length of 75bp.(61) Read pairs were mapped with BOWTIE2 to the indexed version of human genome build USCS hg38 (support.illumina.com/sequencing/sequencing_software/igenome.html) using a local sensitivity mapping algorithm with one allowed base pair mis-match ((64)). The resultant SAM files were converted to BAM format and counted with the Python tool HTSeq.((65)) Each read pair was counted once per annotated genomic feature. The genomic feature count matrix was analyzed using R 4.1.2 (66) and Bioconductor 3.14 (67). Filtering and normalization were performed using the R package edgeR (68). The complete count table was filtered to retain genes with at least 10 reads (lcpm > 0.6) in at least 15 samples resulting in 14,287 retained genes. The filtered counts were normalized using the trimmed mean of M-values normalization procedure. Differentially expressed genes were computed using limma-voom (69). For each time points we computed the Toplists (differentially expressed genes) for the following contrasts: “(BTM 10μm and BTM 1μM) vs. DMSO”. The final output of limma-voom is a table of Log2 fold change, p-values for the defined contrasts tested, as well as Benjamini-Hochberg corrected false discovery p-values (FDR) (70). Volcano plots were generated in R using EnhancedVolcano. For gene set enrichment analysis (GSEA) the log2-FC between BTM treated (10μm and 1μM) versus DMSO treated samples were computed for all 14,287 genes using limma-voom for each time point. These 14,287 log2-FC values were subsequently analyzed with the Broad GSEA tool using GSEA (71)-preranked with the permutation type set to Gene set (1,000 permutations). The gene sets tested for enrichment were from MSigDB.v7.5.1 (C2-canonical pathways gene sets). Gene sets smaller than 15 genes or larger than 300 genes were filtered out, resulting in 1762 gene sets that were tested for each ranked gene list. (All data and analysis can be found in Supplementary Table S6: file RNASeq_BTM3528_Data and Analysis)

##### RNA Extraction and QPCR

For tissue isolated from xenograft models, the tumor was excised, weighed and placed into RNALater (ThermoFisher) and frozen. Thawed tissue was homogenized using metal beads. RNA was extracted using QIAGEN RNeasy midi kit according to the instructions. RNA was reverse transcribed to cDNA (Applied Biosystems – High-Capacity RNA-to-cDNA kit). qPCR was carried out using Applied Biosystems TaqMan™ Fast Advanced Master Mix with Taqman gene expression assays on an ABI 7900HT Real Time PCR machine using a standard protocol.

#### Statistical Analysis of gene expression in Cell Line Screening

The cell line screen utilized 406 cell lines, comprised of 73 hematopoietic tumors and 333 solid tumors. Among the 406 cell lines, 311 have gene expression data available in the CCLE, 308 have gene copy number data, and 286 have mutation status detected in 1561 genes. For hematopoietic tumor cell lines, 57, 56 and 54 have expression, copy number and mutation data respectively. For solid tumor cell lines, 254, 252 and 232 have expression, copy number and mutation data respectively.

#### Gene Expression analysis for biomarker discovery

In the following analysis, tumor cell lines were analyzed collectively to determine whether any genotype or gene expression profiles associated with response to compound. Gene expression data for 14332 genes from Cancer Cell Line Encyclopedia (CCLE) and Dosage response data provided by Crown Bio for 311 cell lines response to BTM-3528 were used in the analysis. The data were first filtered to remove genes with <5 expression level in 90% of the lines. A joins of CCLE epigenetic profiles and dosage response data along gene symbol name is performed, resulting in a dataset of 284 cell lines as samples with 13558 gene expression scores as features for each sample. Each cell line is assigned an area under the curve (AUC) score used as a label target. A regression analysis is then performed to create a model for predicting predict AUC of a cell line with AUC value as response variable and gene expression data as explanatory variable”. The cells lines were split into a training set of 241 cell lines and 43 cell line held as a test dataset. During regression analysis K-fold cross validation on the training dataset was performed with K=5, resulting in 4 groups of 48 cell lines and one group of 49 cell lines. Each flavor of regression experiment is run 5 times on the training dataset, while excluding one of these groups as validation data). Regression utilizing flavors of the generalized linear model (GLM), with tweedie distributions were performed. Four potential target distributions were considered in the model: normal, poisson, gamma and inverse gaussian. The penalty term weight was set at alphas=1,16,64,128. The most effective model was selected as having the lowest mean square error on the validation set and was found to the GLM with normal distribution and alpha=64.

Using the selected regression model, the correlation coefficient of the GLM link function was used to rank gene enrichment features according to their contribution. Gene features were ranked by highest absolute correlation coefficient and select the top N highest values as key features. Each iteration of the K-Fold cross validation experiments provided different gene importance rankings. The overlap between rankings is provided in Figure S7. For each N we selected genes that were found within all 5 iterations of the cross-validation experiment and assemble N rank genelists.

#### Pharmacokinetic Analysis of BTM-3566 in mouse blood

Blood was collected from a tail vein snip into K2-EDTA tubes. Plasma was isolated and a 20 mL sample was protein precipitated with 200 mL of acetonitrile containing 100 ng/mL diclofenac, tolbutamide and labetalol as internal standards. The mixture was vortex-mixed and centrifuged at 13000 rpm for 15 min, 4 °C. An 80 mL aliquot of the supernatant was transferred to a sample plate and mixed with 80 μL water, then the plate was shaken at 800 rpm for 10 min. A 1 mL aliquot was injected on to a Waters ACQUITY UPLC BEH C18 2.1*50mm, 1.7μm reverse phase column using a two-component mobile phase gradient. Mobile phase A was 0.1% trifluoracetic acid in water and mobile phase B 0.1% TFA in acetonitrile. Plasma BTM-3566 was detected using electrospray ionization and multiple reaction monitoring mass spectrometry (BTM-3528 [M+H]+ m/z 522.10>203.1. All data were analyzed using single compartment analysis in WinNonLin.

#### Illustrations

Artwork has been created with Biorender (www.biorender.com) under a standard academic license.

---

**Figure S1.**
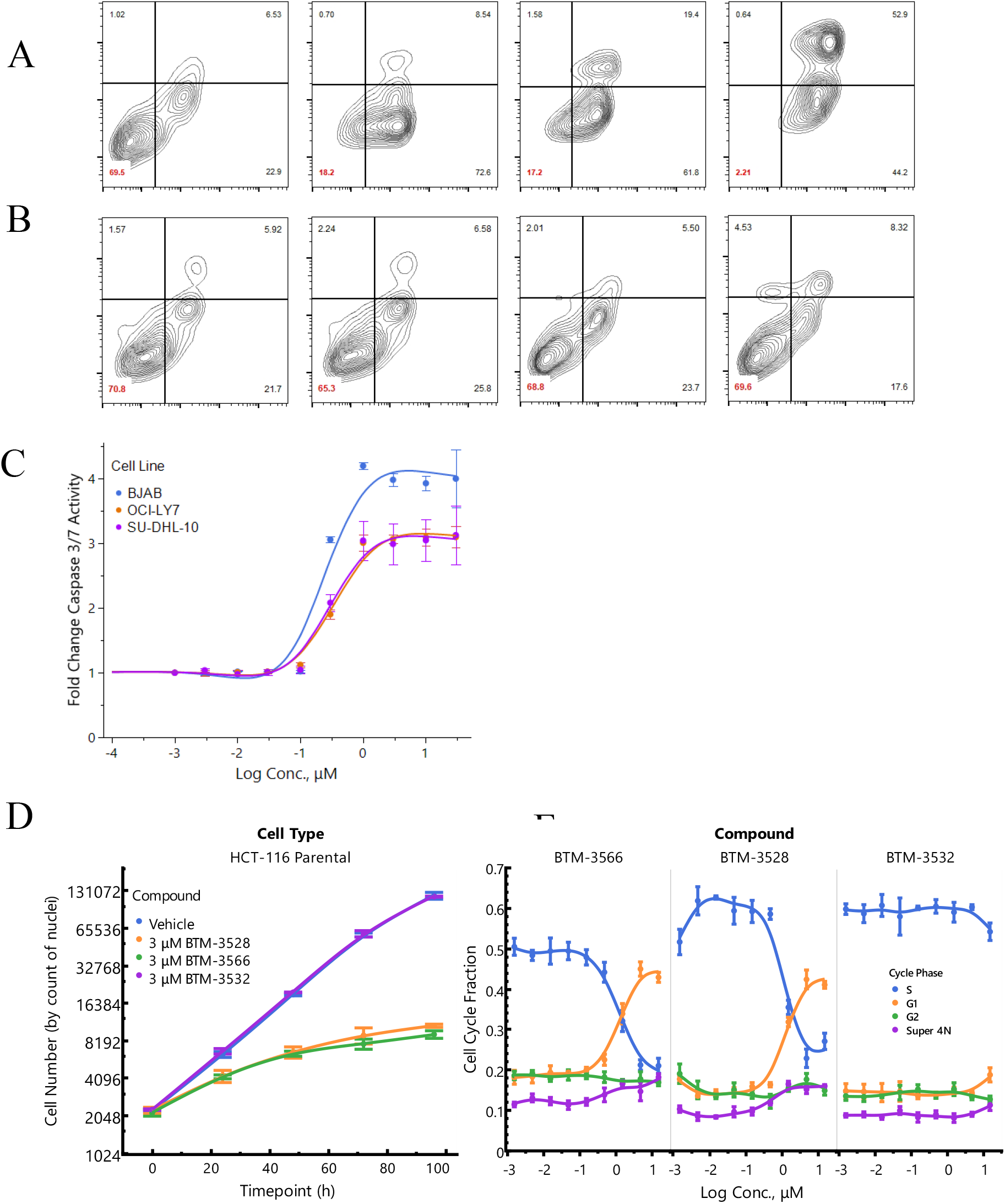
BTM-compounds −3528 and −3566 induce apoptosis through caspase 3/7 activation in DLBCL and G1 cell cycle arrest in solid tumors. **A)** Annexin-PI staining performed in BJAB cells incubated with **2μM BTM3566** for 12,24, and 36 hrs. The percentage of surviving cells is indicated in the lower left quadrant. **B)** Annexin-PI staining performed in BJAB cells incubated with **2μM BTM3532** for 12,24, and 36 hrs. **C)** Caspase 3/7 activity measured by Caspase-Glo assay. Cells were plated in duplicate and incubated for 24 hours then treated with a serial-dilution of **BTM-3566** for 72-hours. Data represent the average fluorescence intensity seen in duplicate experiments. **D)** Growth inhibition by BTM-3528 and BTM-3566 but not BTM-3532. HCT 116 cells were plated into black-walled poly-L-Lysine coated microclear plates and treated with 3 ◻M compound for 24 – 96 hours after which cell number was determined by imaging and counting of DAPI specific nuclear label. **E)** HCT 116 cells were treated with compound and measured for DNA content, DNA synthesis and levels of phosphorylated histone H3.

**Figure S2.**
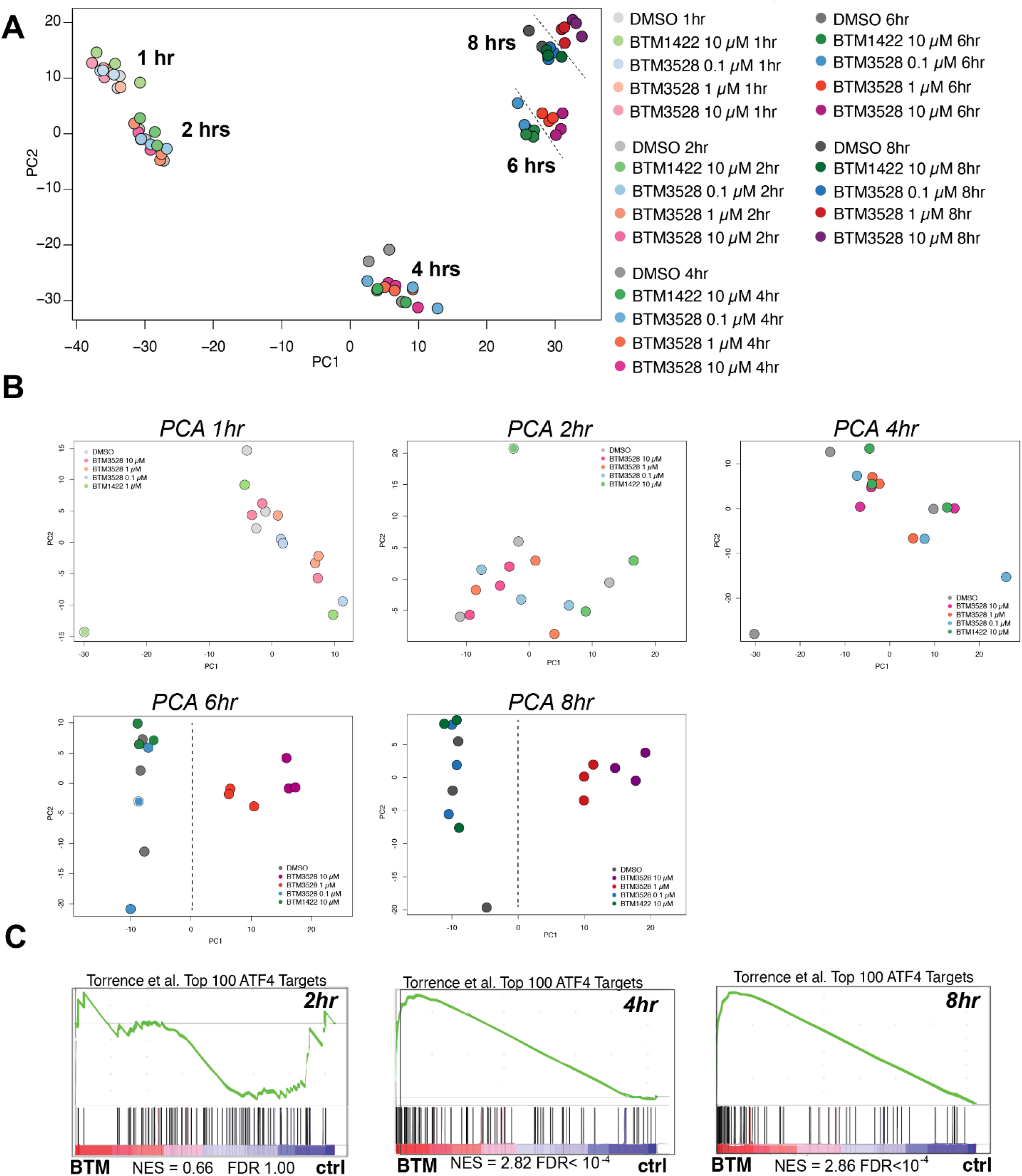
RNASeq of BTM-3566 treated HCT116 cells. **A)** PCA (Principle Component Analysis) of the complete quantile-normalized dataset **B)** PCAs of individual time points (batches) **C)** GSEA-Plot of ATF4 target genes in HCT116 cells treated with 1 μM BTM-3528 versus vehicle-treated HCT116 at the 2, 4 and 8 hour time points.

**Figure S3.**
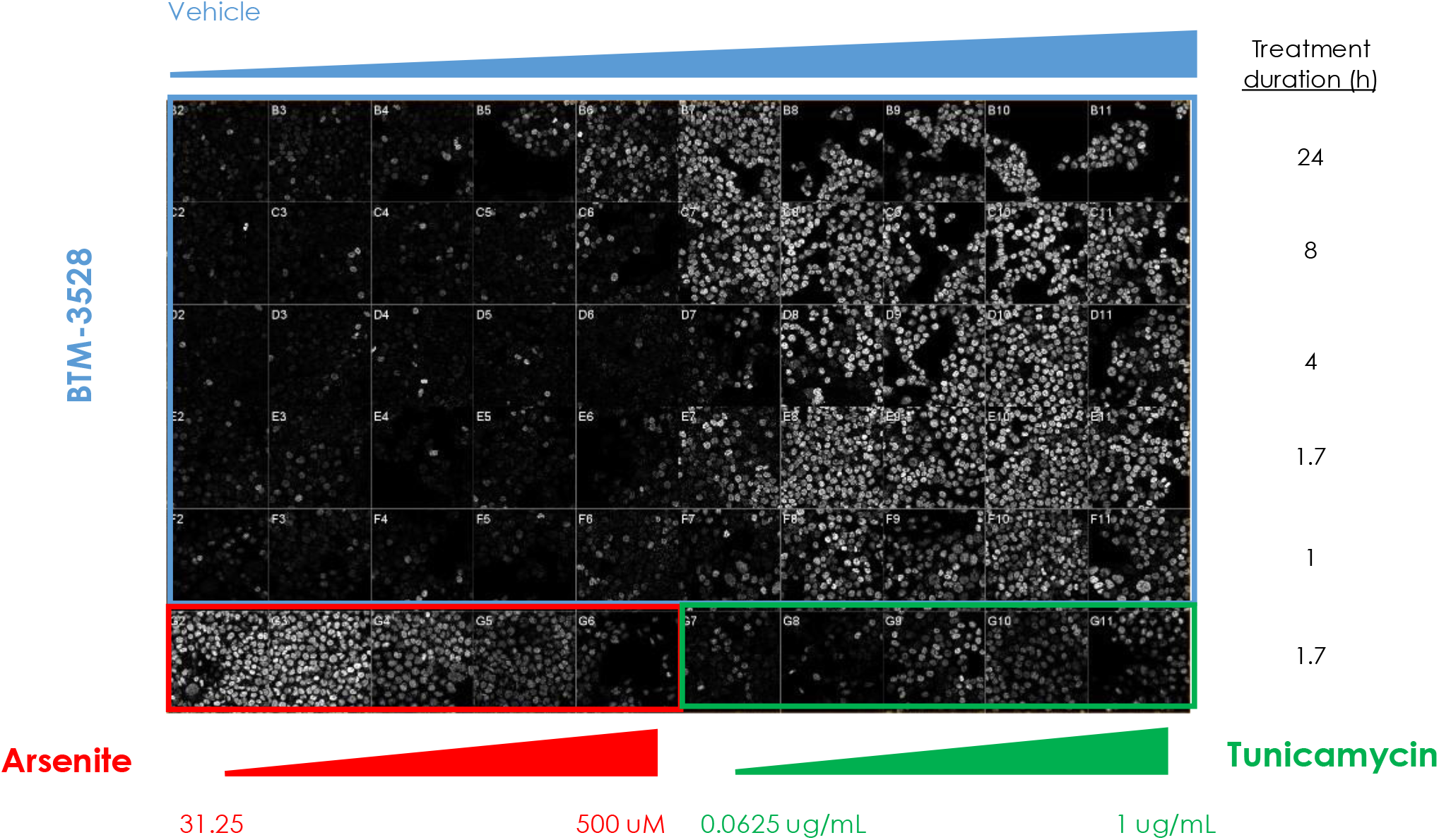
BTM-3528 induces nuclear ATF4 protein accumulation in a dose-dependent manner. HCT-116 cells were plated in 96-well plates and treated with BTM-3528, fixed and ATF4 detected using immunofluorescent imaging. Cell level data were capture for fluorescent intensity of staining for ATF4. Segmentation of nuclear staining was determined with DAPI and nuclear associated ATF4 signal quantified. Mean nuclear staining intensity for each cell was determined in each well. The data are plotted as the median of the mean nuclear intensity in each well. All data are plotted as the mean +/− SD for 3 replicate plates.

**Figure S4.**
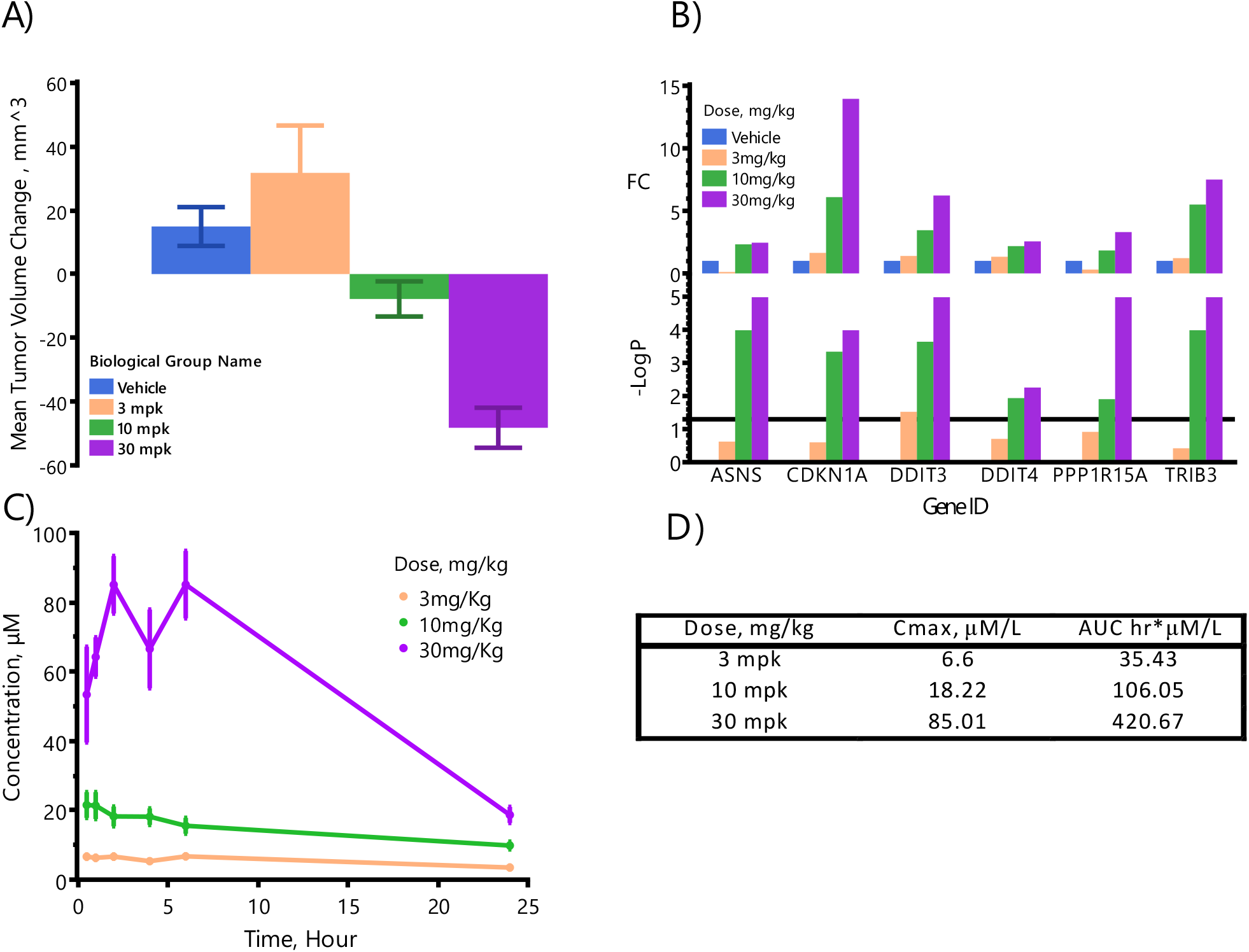
Induction of the ATF4 ISR is correlated with reduction in tumor volume. BTM-3566 was tested in a model of DLBCL in scid mice. SU-DHL-10 cells were used for subcutaneous tumor implantation. Mice were randomized for treatment when tumor volumes reached 200 mm3. BTM-3566 was dosed po, qd. **A)** Changes in tumor volume on day 5 (n=8 +/− SEM); **B)** BTM-3566 induces tumor ATF4 gene expression in a dose dependent manner. On day 5 tumor volumes were recorded, and tissue harvested for QPCR analysis of mRNA expression. Analysis performed with QuantStudio software using three internal reference genes (RPS11, RPSL11, RPL35A) for data normalization. All data are expressed as the composite fold-change (upper panel) and -LogP (lower panel) for significance testing for each gene in each group relative to untreated controls. **C)** Pharmacokinetics of BTM-3566 in mice. On day 1, blood plasma levels of BTM-3566 were determined using LC-MS. **D)** PK parameters following a single dose of BTM-3566 in this experiment.

**Figure S5.**
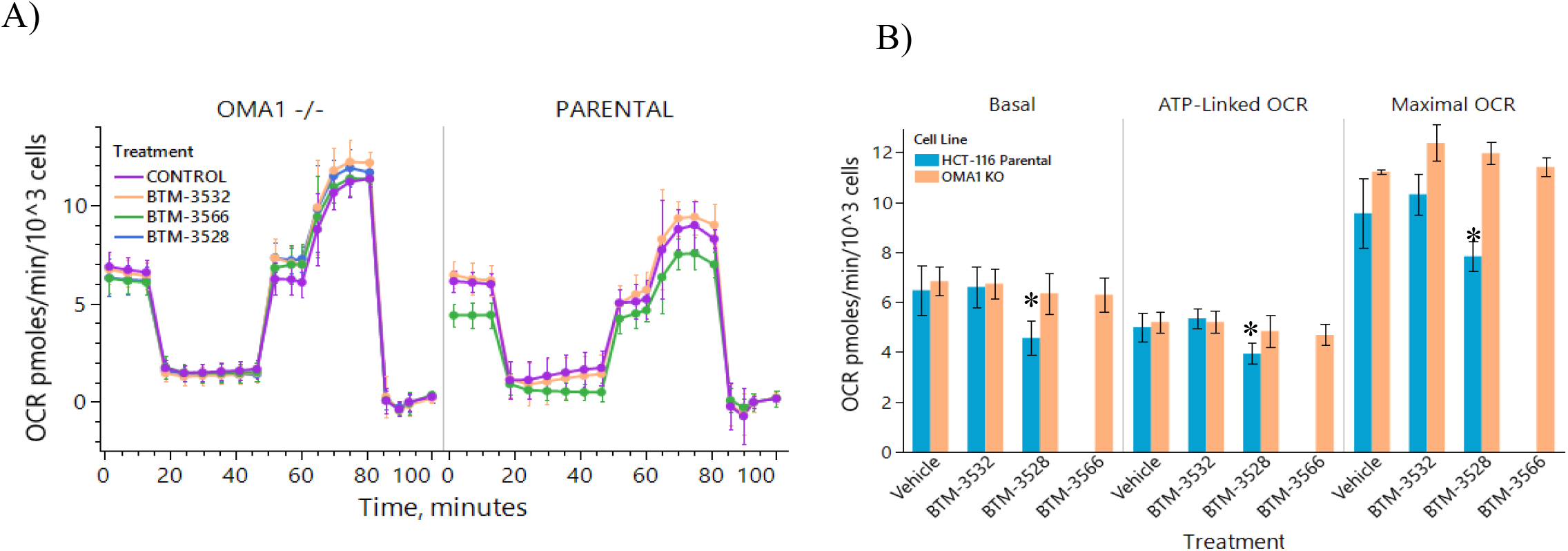
OMA1 is required for BTM mediated reduction in mitochondrial respiration. **A)** HCT-116 Parental or OMA1 −/− cells were treated with 3 mM BTM compounds for 4 hours and then oxygen consumption measured in an Agilent Seahorse device. All data are plotted as the mean +/− SD (n=3) **B)** Basal, ATP linked, maximal OCR were computed for each treatment in each cell line. All data are plotted as the mean +/− SD (n=3 replicates. *, p<.01

**Figure S6.**
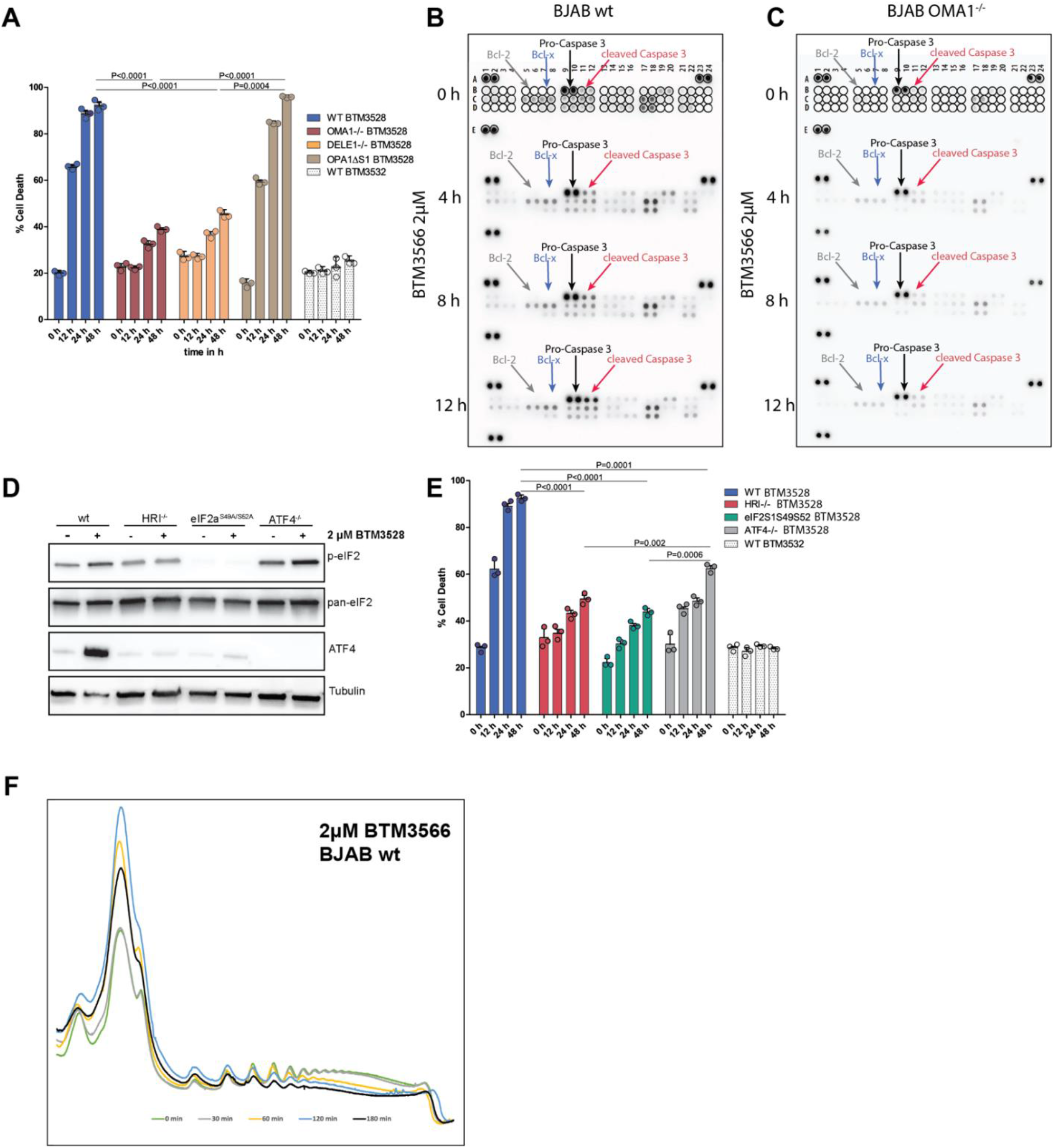
BTM compounds induce the ATF4 ISR via the OMA1-DELE1 mitochondrial quality control pathway Cell Death in parental and OMA1 −/−; DELE1 −/−; and OPA1DS1/DS1 genetically modified BJAB cells. **A)** Cell Death in parental and OMA1^−/−^; DELE1^−/−^ and OPA1*^DS1/DS1^* BJAB cells following treatment with 2 μM BTM-3528. Annexin-Propidium Iodide staining of cells at 12, 24 and 48 hours of exposure. All data are plotted as the mean +/− SD (n=3). **B,C)** Proteome Profiler™ Apoptosis Arrays of BJAB wt and BJAB OMA1^−/−^ cells treated with BTM3566 for 4h,8h, and 12h. **D)** Western Blot analysis of phospho-eIF2a and ATF4 levels in BJAB parental, HRI^−/−^; eIF2a^Ser49/Ser52^ and ATF4 ^−/−^ BJAB cells following 4 hours of treatment with vehicle or 2 μM BTM-3528. **E)** Cell Death in parental, HRI^−/−^; eIF2a^Ser49/Ser52^ and ATF4 ^−/−^ BJAB cells following treatment with 2 μM BTM-3566. Annexin-Propidium Iodide staining of cells at 12, 24 and 48 hours of exposure. All data are plotted as the mean +/− SD (n=3). **F)** Kinetics of Polysome inhibition in BJAB wt cells treated with BTM3566 for 30,60,120, and 180 min.

**Figure S7.**
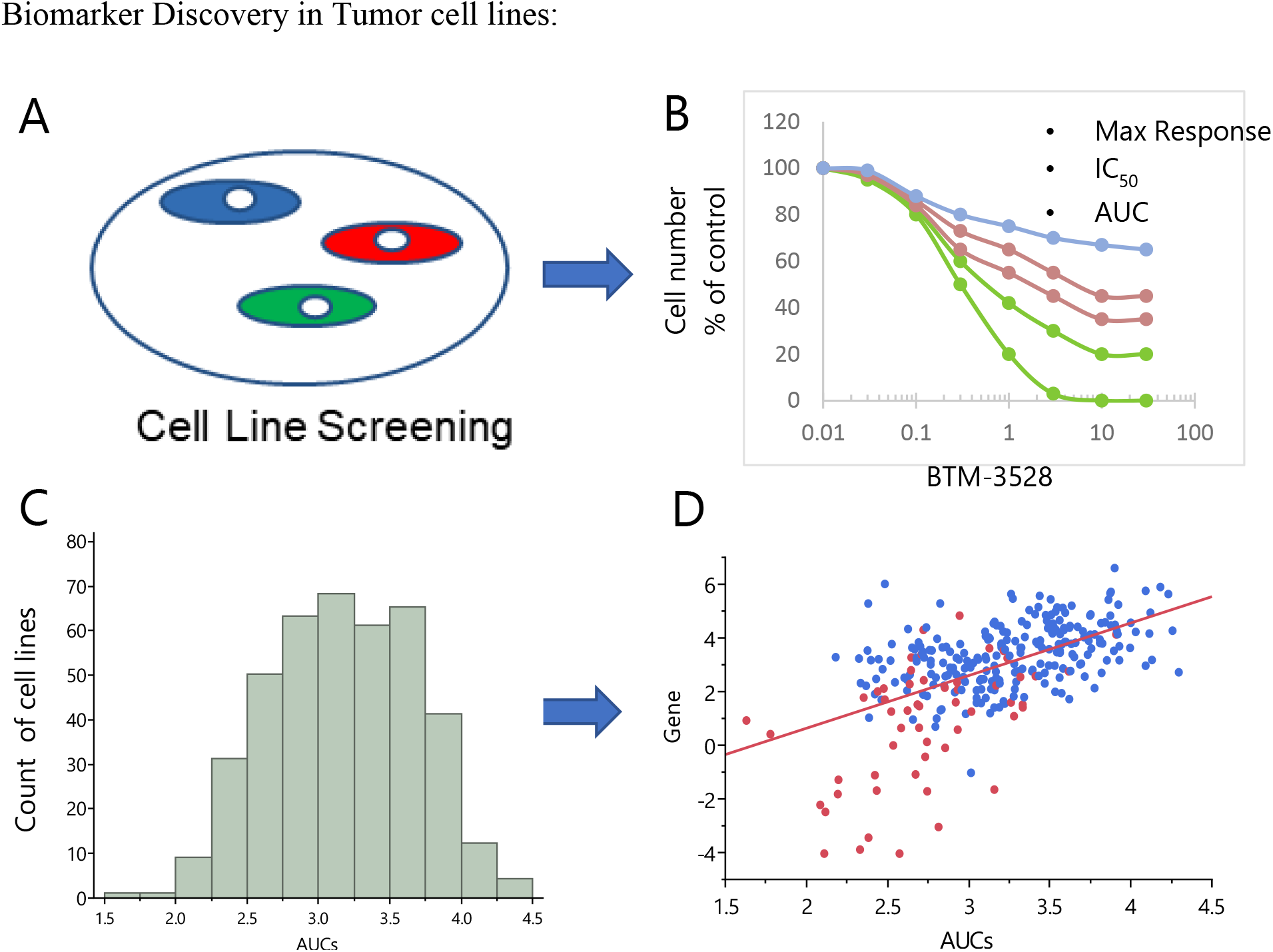
Workflow for analyzing the relationship of global gene expression to response of 406 cell lines to BTM-3528. **A)** 406 cell lines were tested in the analysis. All cells were prepared in appropriate media containing 10-15% fetal bovine serum. BTM-3528 was prepared as a solution in DMSO and diluted into DMSO to provide a final starting concentration of 30 mM. An eight point 3-fold serial dilution of BTM-3528 was prepared in DMSO, mixed into media + FBS and then added to the cells. The final DMSO concentration was 0.1%. The cells were incubated for 96 hours with compound. At the end of the incubation period cell number was determined using Cell-Titre Glo. **B)** Dose response curves were generated for each cell line. Maximum response, IC_50_ and Area under curve (AUC) were determined. In this analysis, the smaller the AUC the more inhibition of cell number. **C)** The distribution frequency of AUC for the cell lines were normally distributed. **D)** CCLE data for gene expression was available for 284 cell lines. Linear regression was performed, and a Spearmans regression score computed.

**Figure S8.**
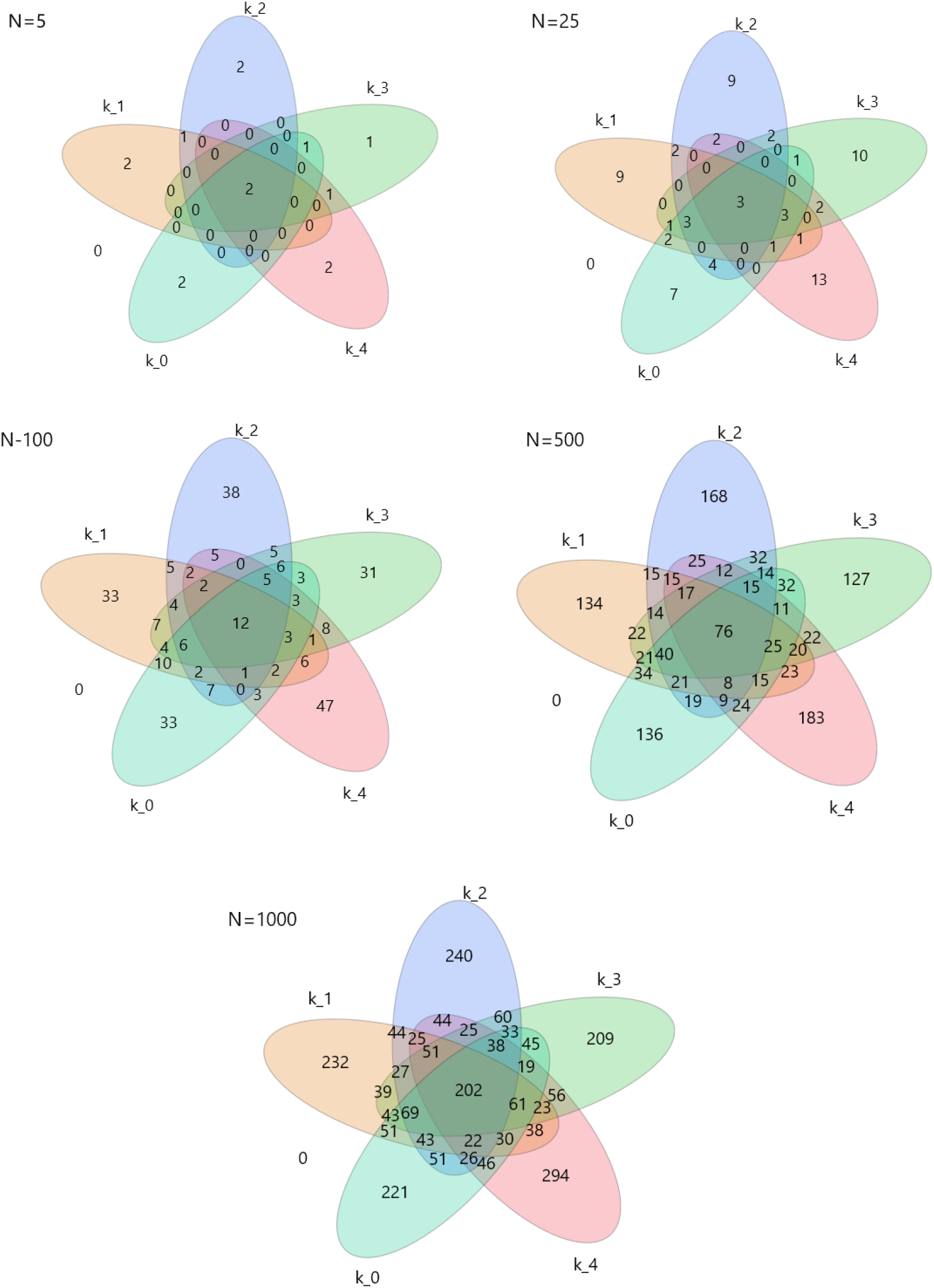
Genes correlating with BTM-3528 activity identified using Using k-fold cross validation. The selected regression model was used to rank gene enrichment features according to contribution. The gene features are ranked by highest absolute corrcoef and the top N highest values are selected as key features. Each iteration of the K-Fold cross validation analysis provided different gene importance rankings. Common genes were visualized between rankings. For each N we selected genes that were found within all 5 iterations of the K-fold cross validation and assembled N_rank genelists as Venn Diagrams. Tabular listing of the genes can be found in Supplemental Table 4_Crown Screening data analysis_Venn Diagram.

**Figure S9.**
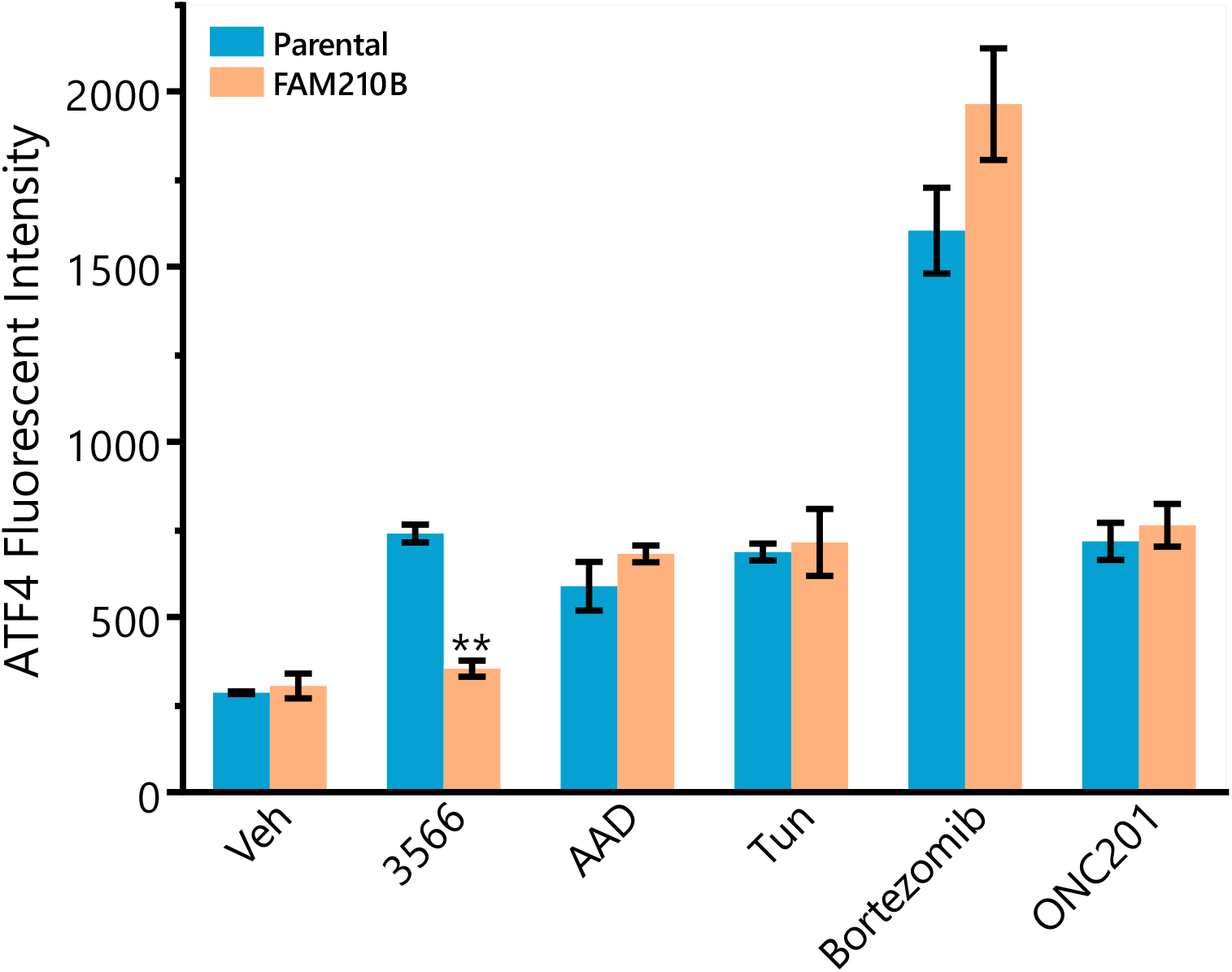
FAM210B regulates BTM-mediated ATF4 protein expression in HCT-116 cells. Parental or FAM210B expressing HCT-116 cells were plated in 96-well plates and treated for 4 hours with 3 μM BTM-3566, 20 nM Bortezomib, 15 μM ONC201 or cultured in culture media deficient in methionine. ATF4 protein was detected using immunofluorescent imaging. Cell level data were capture for fluorescent intensity of staining for ATF4. Segmentation of nuclear staining was determined with DAPI and nuclear associated ATF4 signal quantified. Mean nuclear staining intensity for each cell was determined in each well. The data are plotted as the median of the mean nuclear intensity in each well. All data are plotted as the mean +/− SD for 3 replicate plates. All treatments conditions compared the parental vs the FAM210B expressing cells; ** p<.005

**Figure S10.**
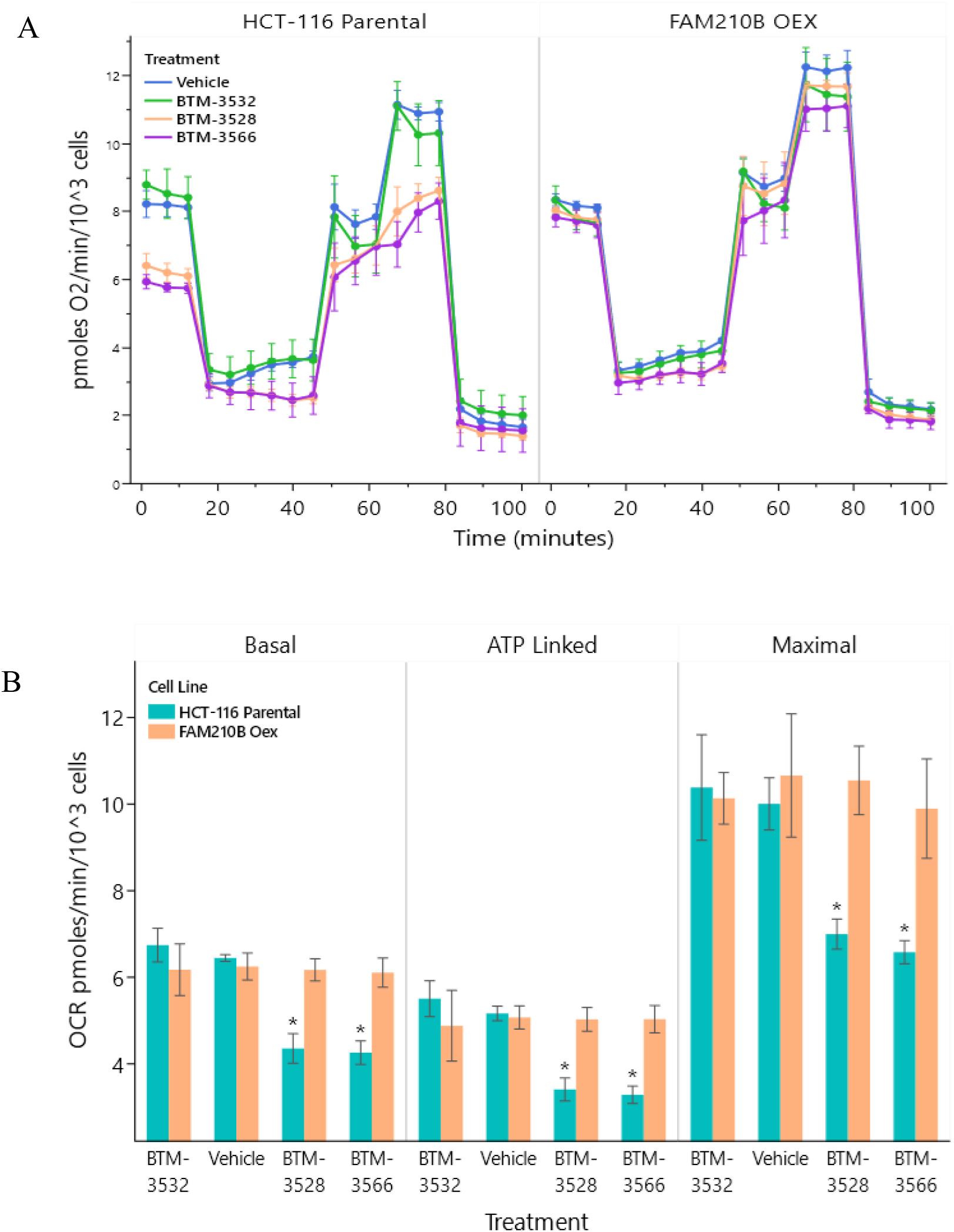
FAM210B overexpression inhibits changes in compound mediated respiration. **A)** Parental or FAM210B stably expressing HCT-116 cells were treated with 3 mM BTM for 4 hours and then oxygen consumption measured in an Agilent Seahorse device. All data are plotted as the mean +/− SD (n=3) **B)** Basal, ATP linked, maximal OCR were computed for each treatment in each cell line. All data are plotted as the mean +/− SD (n=3 replicates. *, p<.05

