## Supplementary material for "Targeting aggressive B-cell lymphomas through pharmacological activation of the mitochondrial protease OMA1": RNA Seq Data: RNASeq_BTM3528_limmaVoom.R

```
################################################################################################
################################################################################################
#### Analysis of BTM 3528 RNASeq in HCT116 cells ###############################################
################################################################################################
################################################################################################
library(edgeR)
library(EnhancedVolcano)
library(limma)
library(UsingR)
library(sva)
library(genefilter)
library(Rtsne)
library(RColorBrewer)
library(gplots)
library(ggplot2)
library(ggsci)
library(rgl)
library(biomaRt)
library(BiocParallel)
bpparam <- MulticoreParam(workers = 10)
bpparam

display.brewer.pal(9, "Blues")
brewer.pal(9, "Blues")

hmcol <- colorRampPalette(brewer.pal(11, "RdBu"))(256)
hmcol <- rev(hmcol)

mypal = pal_npg("nrc", alpha = 7)(9)
mypal

library("scales")
show_col(pal_npg("nrc")(10))
show_col(pal_aaas("default")(10))


################################################################################################
#### 1. Read in data & raw counts ##############################################################
################################################################################################
pData       <- read.delim("RNASeq_BTM3528_Targets.txt",row.names=1,header=TRUE,sep="\t",stringsAsFactors =FALSE)
pData$group <- as.factor(pData$group)
countData   <- read.delim("Serum-Merged-Counts_Scitovation.txt",header=TRUE,sep="\t",stringsAsFactors =FALSE, row.names = 1)       # all 39428 probes 
annotation  <- read.delim("Annotation_RNASeq_BTM3528.txt",row.names=1,header=TRUE,sep="\t",stringsAsFactors =FALSE)

table(colnames(countData)==row.names(pData))
table(row.names(countData)==row.names(annotation))

################################################################################################
#### 2. create edgeR DGEList object: ###########################################################
################################################################################################
dge.raw  <- DGEList(counts  = countData, 
                    genes   = annotation, 
                    samples = pData)  # must contain a factor "group"

# look for library size distribution
hist(dge.raw$samples$lib.size)
median(dge.raw$samples$lib.size)    # 14.5 mio reads
max(dge.raw$samples$lib.size)       # 20 mio reads
min(dge.raw$samples$lib.size)       # 9 mio reads ==> max/min < 3 ==> limma-trend would be possible

# density of log-cpm in raw, unfiltered data
lcpm.dge.raw <- cpm(dge.raw, log=TRUE, prior.count = 1)
simple.densityplot(as.data.frame(lcpm.dge.raw))   
boxplot(lcpm.dge.raw,col=pData$Color,cex.axis=0.5,las=2)

################################################################################################
#### 3. Pre-filtering and TMM-Normalization ####################################################
################################################################################################

#### 3.1 Filter out genes with no or very low counts ###########################################
table(dge.raw$samples$group)  # minimal group size = 15
keep.exprs <- filterByExpr(dge.raw, group=dge.raw$samples$group)  # keeps genes with >=10 reads in [min(groupSize)]= 15 samples = 10/15 Mio = 0.6 cpm   
dge.TMM    <- dge.raw[keep.exprs,,keep.lib.sizes=FALSE]           # recalculates lib size after filtering

lcpm.filter <- cpm(dge.TMM, log=TRUE, prior.count = 1)
simple.densityplot(as.data.frame(lcpm.filter))
boxplot(lcpm.filter,col=pData$Color,cex.axis=0.5,las=2)

#### 3.2 TMM- Normalization ####################################################################

# TMM normalizes the library size, rather then the counts for sequencing depth and RNA composition
# effective library size = library size * norm.factors
dge.TMM <- calcNormFactors(dge.TMM, method = "TMM") # stores norm.factors in $samples$norm.factors
dge.TMM$samples$norm.factors                        # pretty close together ==> TMM only has mild effects in this dataset

eset <- cpm(dge.TMM, log=TRUE, prior.count = 1)     # takes stored norm.factors into account automatically 
simple.densityplot(as.data.frame(eset))
boxplot(eset,col=pData$Color,cex.axis=0.5,las=2)

################################################################################################
#### 4. PCA ####################################################################################
################################################################################################

pca.eset <- prcomp(t(eset))           
plot(pca.eset$x, pch=16, col=pData$Color, cex=1.8,asp=1)
legend(22,-3, legend=unique(pData$Treatment_Time), col=unique(pData$Color), pch=16, bty="n", cex=0.7)

#### overlay expression values of genes over PCA
row.names(pData) == colnames(eset)
pData      <- cbind(as.data.frame(pca.eset$x[,c(1:2)]),pData)
pData$ATF4 <- eset["ATF4",]

ggplot(pData,aes(x=PC1,y=PC2,color=ATF4)) + geom_point(size=4) + 
  scale_shape_manual(values=c(15,1,16,19,17)) +
  theme_bw(base_size = 12, base_family = "") +
  theme(panel.grid.major = element_blank(), panel.grid.minor = element_blank()) +
  scale_colour_gradientn(colours = cbPalette, guide = "colourbar")


################################################################################################
#### 5. Batch correction   #####################################################################
################################################################################################

#### 5.1 Combat ################################################################################
################################################################################################
batch           <- pData$Time.in.hr                          
modcombat       <- model.matrix(~1, data=pData)         
eset.batch      <- ComBat(dat=eset, batch=batch, mod=modcombat,par.prior=TRUE, prior.plots=TRUE)
dev.off()
write.table(eset.batch, file = "ESET_BTM3528_Batchcorrected.txt", sep="\t",row.names = TRUE, col.names = NA)

pca.eset.batch <- prcomp(t(eset.batch))           
plot(pca.eset.batch$x, pch=16, col=pData$Color, cex=2, asp=1)
legend(-28,17, legend=unique(pData$Treatment_Time), col=unique(pData$Color), pch=16, bty="n", cex=0.7)

################################################################################################
#### 6. Subsampling   ##########################################################################
################################################################################################
pData.8hr <- subset(pData, pData$Time.in.hr == 8)
pData.6hr <- subset(pData, pData$Time.in.hr == 6)
pData.4hr <- subset(pData, pData$Time.in.hr == 4)
pData.2hr <- subset(pData, pData$Time.in.hr == 2)
pData.1hr <- subset(pData, pData$Time.in.hr == 1)

dge.TMM.8hr <- dge.TMM[,row.names(pData.8hr)]
dge.TMM.6hr <- dge.TMM[,row.names(pData.6hr)]
dge.TMM.4hr <- dge.TMM[,row.names(pData.4hr)]
dge.TMM.2hr <- dge.TMM[,row.names(pData.2hr)]
dge.TMM.1hr <- dge.TMM[,row.names(pData.1hr)]

eset.8hr <- cpm(dge.TMM.8hr, log=TRUE, prior.count = 1) 
eset.6hr <- cpm(dge.TMM.6hr, log=TRUE, prior.count = 1) 
eset.4hr <- cpm(dge.TMM.4hr, log=TRUE, prior.count = 1) 
eset.2hr <- cpm(dge.TMM.2hr, log=TRUE, prior.count = 1) 
eset.1hr <- cpm(dge.TMM.1hr, log=TRUE, prior.count = 1) 

pca.eset.8hr <- prcomp(t(eset.8hr))           
plot(pca.eset.8hr$x, pch=16, col=pData.8hr$Color, cex=2.5, asp=1)
legend(14,-12, legend=unique(pData.8hr$Treatment_Time), col=unique(pData.8hr$Color), pch=16, bty="n", cex=1)
write.table(eset.8hr,file="ESET_logCPM_TMM_8hrs.txt",sep="\t",col.names=NA)

pca.eset.6hr <- prcomp(t(eset.6hr))           
plot(pca.eset.6hr$x, pch=16, col=pData.6hr$Color, cex=2.5, asp=1)
legend(13,-12, legend=unique(pData.6hr$Treatment_Time), col=unique(pData.6hr$Color), pch=16, bty="n", cex=1)
write.table(eset.6hr,file="ESET_logCPM_TMM_6hrs.txt",sep="\t",col.names=NA)

pca.eset.4hr <- prcomp(t(eset.4hr))           
plot(pca.eset.4hr$x, pch=16, col=pData.4hr$Color, cex=2.5, asp=1)
legend(12,-18, legend=unique(pData.4hr$Treatment_Time), col=unique(pData.4hr$Color), pch=16, bty="n", cex=1)
write.table(eset.4hr,file="ESET_logCPM_TMM_4hrs.txt",sep="\t",col.names=NA)

pca.eset.2hr <- prcomp(t(eset.2hr))           
plot(pca.eset.2hr$x, pch=16, col=pData.2hr$Color, cex=2.5, asp=1)
legend(13,20, legend=unique(pData.2hr$Treatment_Time), col=unique(pData.2hr$Color), pch=16, bty="n", cex=1)
write.table(eset.2hr,file="ESET_logCPM_TMM_2hrs.txt",sep="\t",col.names=NA)


pca.eset.1hr <- prcomp(t(eset.1hr))           
plot(pca.eset.1hr$x, pch=16, col=pData.1hr$Color, cex=2.5, asp=1)
legend(-30,15, legend=unique(pData.1hr$Treatment_Time), col=unique(pData.1hr$Color), pch=16, bty="n", cex=1)


###############################################################################################
#### 7. limma.voom - differentially expressed genes at 8 hrs ##################################
###############################################################################################

#### 7.1. design matrix #######################################################################
groups.8hr       <- factor(pData.8hr$Group)
design.8hr       <- model.matrix(~0+groups.8hr)
rownames(design.8hr) <- row.names(pData.8hr)
colnames(design.8hr) <- c("DMSO","BTM_10", "BTM_1","BTM_0.1","BTM1422")
design.8hr
#### 7.2. contrast matrix ####################################################################
cont.matrix.8hr <- makeContrasts(c1 = (BTM_10+BTM_1)/2 - DMSO,
                                 c2 = BTM_10 - DMSO, 
                                 c3 = BTM_1  - DMSO,
                                 c4 = BTM1422 - DMSO,
                                 levels=design.8hr)
cont.matrix.8hr
#### 7.3. voom ##############################################################################
voom.8hr <- voom(dge.TMM.8hr, design.8hr, plot=TRUE,normalize.method="none") # voom is applied to filtered and TMM-normalized DGE object, prior.count=0.5

#### 7.4. limma #############################################################################
fit.8hr   <- lmFit(voom.8hr,design.8hr)
fit.8hr   <- contrasts.fit(fit.8hr,contrasts=cont.matrix.8hr)
fit2.8hr  <- eBayes(fit.8hr)

top.1.8hr <- topTable(fit2.8hr, coef=1, n=Inf, adjust.method="BH", sort.by="logFC")
top.1.8hr <- top.1.8hr[order(top.1.8hr$logFC,decreasing=TRUE),]
write.table(top.1.8hr,file="Toplist_8hr_BTM10 and BTM1_vs_DMSO.txt",sep="\t",col.names=NA)

top.2.8hr <- topTable(fit2.8hr, coef=2, n=Inf, adjust.method="BH", sort.by="logFC")
top.2.8hr <- top.2.8hr[order(top.2.8hr$logFC,decreasing=TRUE),]
write.table(top.2.8hr,file="Toplist_8hr_BTM10_vs_DMSO.txt",sep="\t",col.names=NA)

top.3.8hr <- topTable(fit2.8hr, coef=3, n=Inf, adjust.method="BH", sort.by="logFC")
top.3.8hr <- top.3.8hr[order(top.3.8hr$logFC,decreasing=TRUE),]
write.table(top.3.8hr,file="Toplist_8hr_BTM1_vs_DMSO.txt",sep="\t",col.names=NA)

top.4.8hr <- topTable(fit2.8hr, coef=4, n=Inf, adjust.method="BH", sort.by="logFC")
top.4.8hr <- top.4.8hr[order(top.4.8hr$logFC,decreasing=TRUE),]
write.table(top.4.8hr,file="Toplist_8hr_BTM1422_vs_DMSO.txt",sep="\t",col.names=NA)

#### 7.5. Volcano Plot ############################################################################
ATF4.Targets <- c("CHAC1","SLC7A11","SLC6A9","SLC1A4","ASNS","DDIT3","DDIT4","ATF4","SARS","PCK2","ADM2","JDP2",
                  "TRIB3", "TSC22D3","PSAT1","STC2","CARS","FAM129A")

EnhancedVolcano(top.1.8hr,lab = rownames(top.1.8hr),x = 'logFC', y = 'adj.P.Val',
                pointSize = 3.5,
                labSize = 4,
                FCcutoff = 1,
                pCutoff = 1e-03,
                caption = '14287 genes, lfc cutoff=1.0; adj.p-value cutoff= 10e-3',
                xlim = c(-2, 4),
                ylim = c(0, -log10(10e-30)),
                xlab = bquote(~Log[2] ~ "fold change"),
                ylab = bquote(~-Log[10] ~ italic(P.adj)),
                title = "BTM3528 vs ctrl 8 hrs",
                subtitle = NULL)

#### 7.6. annotate tophits with BiomaRt #######################################################
library(biomaRt)
top.1.8hr.sig <- subset(top.1.8hr, abs(top.1.8hr$logFC)>0.5 & top.1.8hr$adj.P.Val<0.01)

ensembl    <- useEnsembl(biomart = "genes", dataset = "hsapiens_gene_ensembl")
filters    <- listFilters(ensembl)      # filters    = input of the BM query
attributes <- listAttributes(ensembl)   # attributes = output of the BM query

# get annotation using gene symbols as input / cave unordered output in the order the results are returned from the Biomart server
BM  <- getBM(attributes = c("hgnc_symbol","ensembl_gene_id","external_gene_name","description",
                            "gene_biotype","go_id","name_1006"),
             filters = 'hgnc_symbol', values = top.1.8hr.sig$Gene, mart = ensembl) 

### merge Annotations
Annotation_FINAL$ProbeID <- row.names(Annotation_FINAL)
df.merge                 <- merge(Annotation_FINAL,BM,by.x = "GeneID_FINAL",by.y = "ensembl_gene_id", all.x=TRUE)   
row.names(df.merge)      <- df.merge$ProbeID
df.merge                 <- df.merge[row.names(Annotation_FINAL),]

table(df.merge$GeneID_FINAL == Annotation_FINAL$GeneID_FINAL)
table(row.names(df.merge)   == row.names(Annotation_FINAL)) 

Annotation_FINAL <- df.merge
rm(df.merge)

table(is.na(Annotation_FINAL$GeneID_FINAL))
table(is.na(Annotation_FINAL$external_gene_name))
table(row.names(Annotation_FINAL) == row.names(Annotation.ID026655))

Annotation_FINAL <- Annotation_FINAL[,-3]
colnames(Annotation_FINAL) <- c("GeneID_FINAL","Source_FINAL", "GeneSymbol_FINAL", "GeneName_FINAL", "GeneType_FINAL")


###############################################################################################
#### 8. limma.voom - differentially expressed genes at 6 hrs ##################################
###############################################################################################

#### 8.1. design matrix #######################################################################
groups.6hr       <- factor(pData.6hr$Group)
design.6hr       <- model.matrix(~0+groups.6hr)
rownames(design.6hr) <- row.names(pData.6hr)
colnames(design.6hr) <- c("DMSO","BTM_10", "BTM_1","BTM_0.1","BTM1422")
design.6hr

#### 8.2. contrast matrix ####################################################################
cont.matrix.6hr <- makeContrasts(c1 = (BTM_10+BTM_1)/2 - DMSO,
                                 c2 = BTM_10 - DMSO, 
                                 c3 = BTM_1  - DMSO,
                                 c4 = BTM1422 - DMSO,
                                 levels=design.6hr)
cont.matrix.6hr

#### 8.3. voom ##############################################################################
voom.6hr <- voom(dge.TMM.6hr, design.6hr, plot=TRUE,normalize.method="none") # voom is applied to filtered and TMM-normalized DGE object, prior.count=0.5

#### 8.4. limma ##############################################################################
fit.6hr   <- lmFit(voom.6hr,design.6hr)
fit.6hr   <- contrasts.fit(fit.6hr,contrasts=cont.matrix.6hr)
fit2.6hr  <- eBayes(fit.6hr)

top.1.6hr <- topTable(fit2.6hr, coef=1, n=Inf, adjust.method="BH", sort.by="logFC")
top.1.6hr <- top.1.6hr[order(top.1.6hr$logFC,decreasing=TRUE),]
write.table(top.1.6hr,file="Toplist_6hr_BTM10 and BTM1_vs_DMSO.txt",sep="\t",col.names=NA)

top.2.6hr <- topTable(fit2.6hr, coef=2, n=Inf, adjust.method="BH", sort.by="logFC")
top.2.6hr <- top.2.6hr[order(top.2.6hr$logFC,decreasing=TRUE),]
write.table(top.2.6hr,file="Toplist_6hr_BTM10_vs_DMSO.txt",sep="\t",col.names=NA)

top.3.6hr <- topTable(fit2.6hr, coef=3, n=Inf, adjust.method="BH", sort.by="logFC")
top.3.6hr <- top.3.6hr[order(top.3.6hr$logFC,decreasing=TRUE),]
write.table(top.3.6hr,file="Toplist_6hr_BTM1_vs_DMSO.txt",sep="\t",col.names=NA)

top.4.6hr <- topTable(fit2.6hr, coef=4, n=Inf, adjust.method="BH", sort.by="logFC")
top.4.6hr <- top.4.6hr[order(top.4.6hr$logFC,decreasing=TRUE),]
write.table(top.4.6hr,file="Toplist_6hr_BTM1422_vs_DMSO.txt",sep="\t",col.names=NA)


#### 8.4. Volcano Plot ############################################################################
EnhancedVolcano(top.1.6hr,lab = rownames(top.1.6hr),x = 'logFC', y = 'adj.P.Val',
                pointSize = 3.5,
                labSize = 4,
                FCcutoff = 1,
                pCutoff = 1e-03,
                caption = '14287 genes, lfc cutoff=1.0; adj.p-value cutoff= 10e-3',
                xlim = c(-2, 4),
                ylim = c(0, -log10(10e-30)),
                xlab = bquote(~Log[2] ~ "fold change"),
                ylab = bquote(~-Log[10] ~ italic(P.adj)),
                title = "BTM3528 vs ctrl 6 hrs",
                subtitle = NULL)


###############################################################################################
#### 9. limma.voom - differentially expressed genes at 4 hrs ##################################
###############################################################################################

#### 9.1. design matrix #######################################################################
groups.4hr       <- factor(pData.4hr$Group)
design.4hr       <- model.matrix(~0+groups.4hr)
rownames(design.4hr) <- row.names(pData.4hr)
colnames(design.4hr) <- c("DMSO","BTM_10", "BTM_1","BTM_0.1","BTM1422")
design.4hr

#### 9.2. contrast matrix ####################################################################
cont.matrix.4hr <- makeContrasts(c1 = (BTM_10+BTM_1)/2 - DMSO,
                                 c2 = BTM_10 - DMSO, 
                                 c3 = BTM_1  - DMSO,
                                 c4 = BTM1422 - DMSO,
                                 levels=design.4hr)
cont.matrix.4hr

#### 9.3. voom ##############################################################################
voom.4hr <- voom(dge.TMM.4hr, design.4hr, plot=TRUE,normalize.method="none") # voom is applied to filtered and TMM-normalized DGE object, prior.count=0.5

#### 9.4. limma ##############################################################################
fit.4hr   <- lmFit(voom.4hr,design.4hr)
fit.4hr   <- contrasts.fit(fit.4hr,contrasts=cont.matrix.4hr)
fit2.4hr  <- eBayes(fit.4hr)

top.1.4hr <- topTable(fit2.4hr, coef=1, n=Inf, adjust.method="BH", sort.by="logFC")
top.1.4hr <- top.1.4hr[order(top.1.4hr$logFC,decreasing=TRUE),]
write.table(top.1.4hr,file="Toplist_4hr_BTM10 and BTM1_vs_DMSO.txt",sep="\t",col.names=NA)

top.2.4hr <- topTable(fit2.4hr, coef=2, n=Inf, adjust.method="BH", sort.by="logFC")
top.2.4hr <- top.2.4hr[order(top.2.4hr$logFC,decreasing=TRUE),]
write.table(top.2.4hr,file="Toplist_4hr_BTM10_vs_DMSO.txt",sep="\t",col.names=NA)

top.3.4hr <- topTable(fit2.4hr, coef=3, n=Inf, adjust.method="BH", sort.by="logFC")
top.3.4hr <- top.3.4hr[order(top.3.4hr$logFC,decreasing=TRUE),]
write.table(top.3.4hr,file="Toplist_4hr_BTM1_vs_DMSO.txt",sep="\t",col.names=NA)

top.4.4hr <- topTable(fit2.4hr, coef=4, n=Inf, adjust.method="BH", sort.by="logFC")
top.4.4hr <- top.4.4hr[order(top.4.4hr$logFC,decreasing=TRUE),]
write.table(top.4.4hr,file="Toplist_4hr_BTM1422_vs_DMSO.txt",sep="\t",col.names=NA)

#### 5.4. Volcano Plot ############################################################################
EnhancedVolcano(top.1.4hr,lab = rownames(top.1.4hr),x = 'logFC', y = 'adj.P.Val',
                pointSize = 3.5,
                labSize = 4,
                FCcutoff = 1,
                pCutoff = 1e-03,
                caption = '14287 genes, lfc cutoff=1.0; adj.p-value cutoff= 10e-3',
                xlim = c(-2, 4),
                ylim = c(0, -log10(10e-30)),
                xlab = bquote(~Log[2] ~ "fold change"),
                ylab = bquote(~-Log[10] ~ italic(P.adj)),
                title = "BTM3528 vs ctrl 4 hrs",
                subtitle = NULL)

###############################################################################################
#### 10. limma.voom - differentially expressed genes at 2 hrs #################################
###############################################################################################

#### 10.1. design matrix #######################################################################
groups.2hr       <- factor(pData.2hr$Group)
design.2hr       <- model.matrix(~0+groups.2hr)
rownames(design.2hr) <- row.names(pData.2hr)
colnames(design.2hr) <- c("DMSO","BTM_10", "BTM_1","BTM_0.1","BTM1422")
design.2hr

#### 10.2. contrast matrix ####################################################################
cont.matrix.2hr <- makeContrasts(c1 = (BTM_10+BTM_1)/2 - DMSO,
                                 c2 = BTM_10 - DMSO, 
                                 c3 = BTM_1  - DMSO,
                                 c4 = BTM1422 - DMSO,
                                 levels=design.2hr)
cont.matrix.2hr

#### 10.3. voom ##############################################################################
voom.2hr <- voom(dge.TMM.2hr, design.2hr, plot=TRUE,normalize.method="none") # voom is applied to filtered and TMM-normalized DGE object, prior.count=0.5

#### 10.4. limma ##############################################################################
fit.2hr   <- lmFit(voom.2hr,design.2hr)
fit.2hr   <- contrasts.fit(fit.2hr,contrasts=cont.matrix.2hr)
fit2.2hr  <- eBayes(fit.2hr)

top.1.2hr <- topTable(fit2.2hr, coef=1, n=Inf, adjust.method="BH", sort.by="logFC")
top.1.2hr <- top.1.2hr[order(top.1.2hr$logFC,decreasing=TRUE),]
write.table(top.1.2hr,file="Toplist_2hr_BTM10 and BTM1_vs_DMSO.txt",sep="\t",col.names=NA)

top.2.2hr <- topTable(fit2.2hr, coef=2, n=Inf, adjust.method="BH", sort.by="logFC")
top.2.2hr <- top.2.2hr[order(top.2.2hr$logFC,decreasing=TRUE),]
write.table(top.2.2hr,file="Toplist_2hr_BTM10_vs_DMSO.txt",sep="\t",col.names=NA)

top.3.2hr <- topTable(fit2.2hr, coef=3, n=Inf, adjust.method="BH", sort.by="logFC")
top.3.2hr <- top.3.2hr[order(top.3.2hr$logFC,decreasing=TRUE),]
write.table(top.3.2hr,file="Toplist_2hr_BTM1_vs_DMSO.txt",sep="\t",col.names=NA)

top.4.2hr <- topTable(fit2.2hr, coef=4, n=Inf, adjust.method="BH", sort.by="logFC")
top.4.2hr <- top.4.2hr[order(top.4.2hr$logFC,decreasing=TRUE),]
write.table(top.4.2hr,file="Toplist_2hr_BTM1422_vs_DMSO.txt",sep="\t",col.names=NA)

#### 10.5. Volcano Plot ############################################################################
EnhancedVolcano(top.1.2hr,lab = rownames(top.1.2hr),x = 'logFC', y = 'adj.P.Val',
                pointSize = 3.5,
                labSize = 4,
                FCcutoff = 1,
                pCutoff = 1e-03,
                caption = '14287 genes, lfc cutoff=1.0; adj.p-value cutoff= 10e-3',
                xlim = c(-2, 4),
                ylim = c(0, -log10(10e-30)),
                xlab = bquote(~Log[2] ~ "fold change"),
                ylab = bquote(~-Log[10] ~ italic(P.adj)),
                title = "BTM3528 vs ctrl 2 hrs",
                subtitle = NULL)


###############################################################################################
#### 11. Global Heatmap & PCA of Toplist ######################################################
###############################################################################################
Topgenes.8hr.up    <- subset(top.1.8hr, top.1.8hr$logFC > 1 & top.1.8hr$adj.P.Val<0.05)
Topgenes.8hr.down  <- subset(top.1.8hr, top.1.8hr$logFC < (-1) & top.1.8hr$adj.P.Val<0.05)
Topgenes.8hr       <- rbind(Topgenes.8hr.up,Topgenes.8hr.down)
pData.ordered      <- pData[c(1:15,61:75,46:60,31:45,16:30),]    # order columns according to time
eset.ordered       <- as.matrix(eset[,row.names(pData.ordered)]) # order columns according to time

eset.top.8hr       <- eset.ordered[row.names(Topgenes.8hr),]
table(row.names(pData.ordered) == colnames(eset.top.8hr))

pca.eset.top.8hr <- prcomp(t(eset.top.8hr))           
plot(pca.eset.top.8hr$x, pch=16, col=pData.ordered$Color, cex=2, asp=1)
legend(4,7.5, legend=unique(pData.ordered$Treatment_Time), col=unique(pData.ordered$Color), pch=16, bty="n", cex=0.6)

heatmap.2(eset.top.8hr, col = hmcol, cexRow=0.7, cexCol=0.6,Colv = F,Rowv = F,
          scale="row", trace="none", ColSideColors=pData.ordered$Color, labRow = row.names(eset.top.8hr), labCol=row.names(pData.ordered) )

#### 9.3 PCA with Top gene expression #########################################################
pData.ordered          <- cbind(as.data.frame(pca.eset.top.8hr$x[,c(1:2)]),pData.ordered)
pData.ordered$ATF4     <- eset.ordered["ATF4",] 
pData.ordered$ATF6     <- eset.ordered["ATF6",] 
pData.ordered$DDIT3    <- eset.ordered["DDIT3",] 
pData.ordered$DDIT4    <- eset.ordered["DDIT4",] 
pData.ordered$ASNS     <- eset.ordered["ASNS",] 
pData.ordered$PPP1R15A <- eset.ordered["PPP1R15A",] 
pData.ordered$SARS     <- eset.ordered["SARS",] 
pData.ordered$SLC7A11  <- eset.ordered["SLC7A11",] 

cbPalette <- colorRampPalette(brewer.pal(11, "RdYlBu"))(20)
cbPalette <- rev(cbPalette)

ggplot(pData.ordered,aes(x=PC1,y=PC2,color=ATF4)) + geom_point(size=4.3) + 
  theme_bw(base_size = 12, base_family = "") +
  theme(panel.grid.major = element_blank(), panel.grid.minor = element_blank()) +
  scale_colour_gradientn(colours = cbPalette, guide = "colourbar") +
  coord_fixed(ratio = 1)


##################################################################################################
#### 8. ssGSEA  #################################################################################
#################################################################################################

#### 8.1. ssGSEA compatible matrix ##############################################################
#################################################################################################
matrix.gsea    <- as.matrix(eset[,row.names(pData.ordered)])
matrix.gsea    <- subset(matrix.gsea, row.names(matrix.gsea) %in% genes)
matrix.gsea    <- matrix.gsea[!duplicated(row.names(matrix.gsea)),]

#### 8.2. GeneSets ##############################################################################
#################################################################################################
library(GSEABase)
sets.ssGSEA <- read.delim("Top30_GeneSets_C2_CP.txt",header=FALSE,sep="\t",stringsAsFactors =FALSE)
## read in complete Broad gmt file into GSEABase 
c2.CP       <- getGmt("c2.cp.v7.5.1.symbols.gmt", collectionType=BroadCollection(category="c2"), geneIdType=SymbolIdentifier())
## subset Top30 gene sets
c2.CP.Top30 <- c2.CP[sets.ssGSEA$V1]

#### 8.3. ssgsea ################################################################################
################################################################################################
library(GSVA)
matrix.zscore <- gsva(matrix.gsea, c2.CP.Top30,
                      method="ssgsea",
                      abs.ranking=FALSE,
                      min.sz=15,
                      max.sz=301,
                      mx.diff=TRUE,
                      verbose=TRUE)

hist(as.vector(matrix.zscore))   # ssGSEA is not centered around 0!! ==> shift for correction

heatmap.2(matrix.zscore, col = hmcol, cexRow=0.7, cexCol=0.6,Colv = F,Rowv = F,
          scale="row", trace="none", ColSideColors=pData.ordered$Color, labRow = NULL, labCol=row.names(pData.ordered) )

### visualize 
pData.ordered$ATF4.Targets <- matrix.zscore["Manning_Top100_ATF4 Target Genes_Tunicamycin",] 

ggplot(pData.ordered,aes(x=PC1,y=PC2,color=ATF4.Targets)) + geom_point(size=4.3) + 
  theme_bw(base_size = 12, base_family = "") +
  theme(panel.grid.major = element_blank(), panel.grid.minor = element_blank()) +
  scale_colour_gradientn(colours = cbPalette, guide = "colourbar") +
  coord_fixed(ratio = 1)


##################################################################################################
#### 9. GSEA Plots  ##############################################################################
##################################################################################################

# uses output from BROAD GSEA Tool on the .gct files created above 

# BTM vs DMSO
GSEA.8hr <- read.delim("GSEA_results_BTM_vs_DMSO_voom_8hrs_FINAL.txt",row.names=1,header=TRUE,sep="\t",stringsAsFactors =FALSE)

plot(log10(GSEA.8hr$FDR),GSEA.8hr$NES, ylab="Normalized enrichment score (NES)",
     xlab="FDR (log10)", col=GSEA.8hr$Color, pch=16, cex=2.7)        
segments(log10(0.05),-5,log10(0.05),5, lty = 2)
legend(-5,1, legend=unique(GSEA.8hr$Farbe), col=unique(GSEA.8hr$Color), pch=16, bty="n", cex=1.2)


##################################################################################################
#### 10. Compare to logFC file from Matthew ######################################################
##################################################################################################

logFC.3528.10uM.8hrs <- subset(RNASeq.data, RNASeq.data$Time == "ht8" &
                                            RNASeq.data$Compound.ID_Dose == "3528_10 _M")

logFC.3528.10uM.8hrs.sig <- subset(logFC.3528.10uM.8hrs, logFC.3528.10uM.8hrs$abs_Log2FC>=1.25)

logFC.3528.10uM.6hrs.sig <- subset(RNASeq.data, RNASeq.data$Time == "ht6" & 
                                                RNASeq.data$Compound.ID_Dose == "3528_10 _M" &
                                                RNASeq.data$abs_Log2FC>=1.25)
                                   
RNASeq.data.Matt.3528        <- subset(RNASeq.data, RNASeq.data$Compound.ID == "3528")
RNASeq.data.Matt.sign        <- subset(RNASeq.data.Matt.3528, RNASeq.data.Matt.3528$abs_Log2FC>=1.25)
RNASeq.data.Matt.sign.unique <- RNASeq.data.Matt.sign[!duplicated(RNASeq.data.Matt.sign$Gene),]

RNA.Seq.data.sig.all <- subset(RNASeq.data, RNASeq.data$abs_Log2FC>=1.25)
```
